# Elucidating the Influence of Linker Histone Variants on Chromatosome Dynamics and Energetics

**DOI:** 10.1101/660076

**Authors:** Dustin C. Woods, Jeff Wereszczynski

## Abstract

Linker histones are epigenetic regulators that bind to nucleosomes and alter chromatin structures and dynamics. Biophysical studies have revealed two binding modes in the linker histone/nucleosome complex, the chromatosome, where the linker histone is either centered on or askew from the dyad axis. Each has been posited to have distinct effects on chromatin, however the molecular and thermodynamic mechanisms that drive them and their dependence on linker histone compositions remain poorly understood. We present molecular dynamics simulations of chromatosomes with the globular domain of two linker histone variants, generic H1 (genGH1) and H1.0 (GH1.0), to determine how their differences influence chromatosome structures, energetics, and dynamics. Results show that both unbound linker histones adopt a single compact conformation. Upon binding, DNA flexibility is reduced, resulting in increased chromatosome compaction. While both variants enthalpically favor on-dyad binding, energetic benefits are significantly higher for GH1.0, suggesting that GH1.0 is more capable than genGH1 of overcoming the large entropic reduction required for on-dyad binding which helps rationalize experiments that have consistently demonstrated GH1.0 in on-dyad states but that show genGH1 in both locations. These simulations highlight the thermodynamic basis for different linker histone binding motifs, and details their physical and chemical effects on chromatosomes.

## Introduction

In eukaryotes, chromosomes serve as the primary storage medium of genomic information within an organism and consist predominantly of organized, long condensed fibers of DNA and structural proteins.^1^ These fibers are made of compacted repeating arrays of DNA-protein complexes collectively known as chro-matin. ^2,3^ Despite being tightly condensed, chromatin still allows for enzyme induced replication, repair, and transcription. ^4–6^ The basic building block of chromatin fibers is the nucleosome core particle (NCP) which is comprised of 147 base pairs of DNA wrapped around an octameric core of histone proteins that are built from duplicates of four histones: H2A, H2B, H3, and H4 ^1,7,8^. These histones bind to one another to form H2A-H2B and H3-H4 dimers, while the H3-H4 dimers associate into a tetramer. This tetramer then combines with the H2A-H2B dimers to form the octameric core. ^9,10^

The chromatosome is an extension of the NCP containing the same structural foundations with an additional ∼20 base pairs of DNA accompanied by a linker histone (LH) protein (Figure 1).^11^ Colloquially known as histone H1, this nuclear protein plays a crucial role in the condensation of nucleosome chains into higher order structures, ^12–15^ as well as other cellular functions ^14^ such as gene expression, ^16,17^ heterochromatin genetic activity, ^18^ and cell differentiation, ^19,20^ among many others. ^21–23^ Additionally, linker histones predominantly interact electrostatically with the backbone phosphates of DNA using positively charged residues, ^24–26^ which stabilizes nucleosome arrays hindering linker DNA accessibility. ^15,27–30^ However, this effect has shown to be completely abrogated upon the addition of nucleosome-free regions within H1-saturated arrays. ^31^ They are found roughly every 200 ± 40 base pairs, ^32^ but may be spaced more intermittently to regulate DNA accessibility for transcription factors.

**Figure 1:**
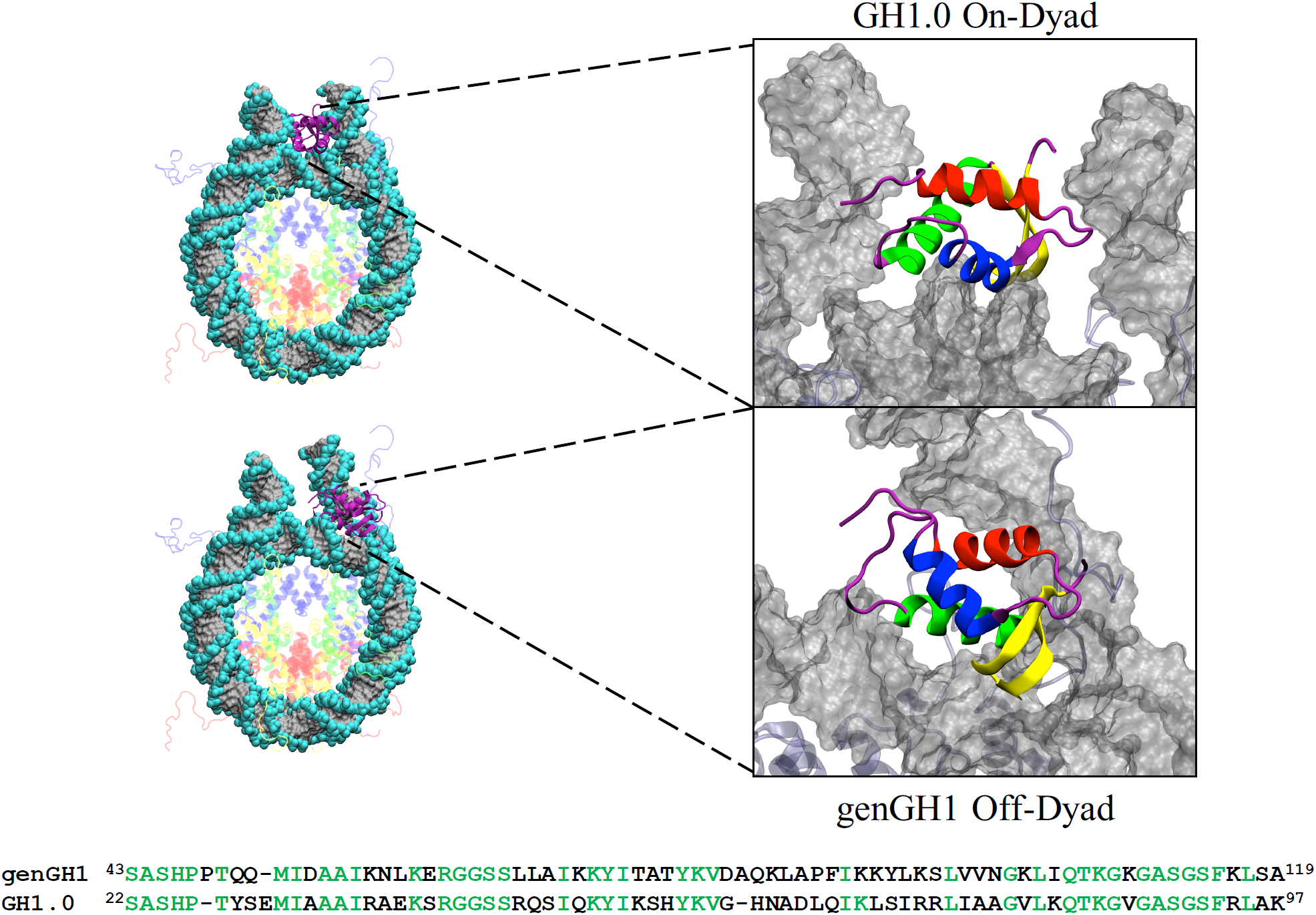
On-dyad (top) and off-dyad (bottom) chromatosome structures. Global structures are shown on the left with zoomed-in figures of the linker histones GH1.0 (top) and genGH1 (bottom) are on the right. On the left, histones are color-coded as follows: Histone H2A (yellow), Histone H2B (red), Histone H3 (blue), Histone H4 (green), Histone genGH1/GH1.0 (purple), and DNA (gray). On the right the linker histones are colored by secondary structure: α-helix 1 (α1; red), α-helix 2 (α2; blue), α-helix 3 (α3; green), the β-sheet (yellow), and disordered regions (purple). Sequences for genGH1 and GH1.0 are shown on the bottom, with identical residues in green.

Linker histones primarily bind to the nucleosome in two states. In the “on-dyad” location ^33–35^ the linker histone is centered on the dyad axis (Figure 1 (top)), whereas in the “off-dyad” configuration the histone binds in a DNA groove off the dyad axis ^36–38^ (Figure 1 (bottom)). The model studied here has the on-dyad linker histone interacting with the DNA minor groove, while in the off-dyad state it is adjacent to the DNA major groove at the +0.5 superhelical location, roughly three to seven base pairs from the dyad axis. Variations in the linker histone binding mode may result in differences in the mechanical stability and overall packaging within the greater chromatin architecture, which would naturally affect the accessibility of DNA in nuclear processes. ^39–41^ This effect was demonstrated in recent coarse-grained simulations by Perišić *et al.* where they found the off-dyad binding mode, as observed in the experimentally generated cryo-EM image of polynucleosomal arrays, ^37^ is a better chromatin condenser than other binding modes. ^41^ Additionally, it was suggested that on-dyad binding is relatively poor for compaction, even compared to systems with hybrid binding modes, suggesting it plays a role in chromatin transcriptional accessibility and dynamic architecture. In a more recent cryo-EM and crystallography study, Garcia-Saez *et al.* showed that the on-dyad state can create alternative compact chromatin conformations, and that shifts in the ionic conditions can induce untwisting to reveal a ladder-like structures. ^42^ Altogether, these works suggest that fluctuating linker histone binding modalities can lead to different levels and structures of chromatin compaction in cooperation with varying linker DNA lengths between nucleosomes. ^43–45^

Several factors contribute to the thermodynamic preference for on-vs off-dyad binding in linker histones. Recent work by Zhou *et al.* examined the binding modes of the wild type linker histone globular domains of *D. melanogaster* generic H1 (genGH1), *G. gallus* H1.0 (GH1.0), and an H1.0 pentamutant. ^36^ Paramagnetic relaxation enhancement (PRE) experiments showed that GH1.0 binds on-dyad and that genGH1 binds off-dyad, but also that a small number of mutations are able to shift the equilibrium of the GH1.0 binding state from on- to off-dyad. ^36^ This suggests that the thermodynamic balance between these states is finely tuned by specific linker histone/nucleosome contacts. These studies are supported by cryo-EM experiments of condensed nucleosome arrays which suggested an off-dyad generic H1 binding mode. ^37^ In contrast, in cryo-EM experiments of single nucleosomes, generic H1 has been observed in the on-dyad binding state, ^34^ while there is evidence to suggest that off-dyad binding in the cryo-EM map of the 30-nm fiber may be a result of cross-linker effects. ^46^ Taken together, these experiments paint the picture that linker histones likely bind in an ensemble of on- and off-dyad states, and that the balance of these two conformations is dictated by several factors including the linker histone primary sequence, the chromatosome’s stereochemical environment, and greater chromatin architecture. ^47^ Note that the linker histone naming convention used here is consistent with that found in the Histone Database and introduced by Talbert *et al.* and may differ from that found in previous studies. ^48,49^

The contrasting influence of linker histone variants on the chromatosome structure and energetics, and the extent to which they affect greater chromatin dynamics, remains unclear. ^50–52^ Using Brownian dynamic docking simulations, Öztürk *et al.* found that GH1.0 displays a range of conformational flexibility and affects the overall chromatosome dynamics, including the linker DNA. ^53^ With similar techniques they later showed that even slightly varied linker histone sequences, including point mutations and posttranslational modifications, can significantly affect the chromatosome structure. ^47,54^ Moreover, with accelerated molecular dynamics simulations they found that the GH1.0 β-sheet loop (β-loop), which has both an open and closed-state in the crystal structure, ^55^ favors the closed-state in solution, although the open-state may still be populated. This is in line with the closed-state being the only conformation observed in chromatosome crystal structures ^33,34,46^. However, there is evidence suggesting that linker histones may exist in alternative conformations ^56,57^ and binding orientations. ^56,58–60^

Despite these and many more ^61^ excellent experimental and computational studies, several questions remain concerning linker histones and their nucleosome binding. For example, to what extent does linker histone plasticity affect its function? What are the effects of on- and off-dyad binding on chromatosome dynamics? How do specific thermodynamic forces influence the on-vs off-dyad binding equilibrium? How are these properties influenced by different linker histone variants? To address these questions, we have performed a series of conventional and free energy molecular dynamics (MD) simulations of chromatosomes containing the *Drosohila melanogaster* generic globular domain of H1 (genGH1) and *Gallus gallus* globular domain of H1.0 (GH1.0, which has previously been referred to as H5) bound in both on- and off-dyad states. Building off work from the Bai and Wade groups, this work was focused on the globular domain of each linker histone.^36,38,47,53,54,56^ Results suggest that both genGH1 and GH1.0 readily adopt a single compact configuration. Furthermore, in the off-dyad state linker histones display increased localized sampling while modestly altering the linker DNA dynamics, while linker histones have highly stable binding in the on-dyad state while significantly restricting DNA motions. Energetic analyses shows that the equilibrium between on- and off-dyad binding is the result of a balance between Van der Waals and electrostatic interactions that is dictated by the linker histone variant type. Together, these results suggest that, regardless of the variant, on-dyad binding is enthalpically stabilized whereas off-dyad binding is relatively more entropically stabilized. Furthermore, when in on- and off-dyad conformations, different linker histone variants have similar effects on chromatosome structures and dynamics, and that the role of linker histone modifications is likely to shift the relative populations between these binding states. The *in vitro* ensemble of binding modes, and therefore the greater structure of linker histone containing chromatin fibers, is therefore dictated by competing thermodynamic forces which are likely influenced by a myriad of structural and environmental factors *in vivo*.

## Methods

### System construction

Core histones were modelled based on the 1KX5 crystal structure (resolution 1.94 Å ^62^). The asymmetric Widom 601 DNA ^63,64^ was taken from the 4QLC crystal structure, which has a lower resolution (3.50 Å) but both DNA and GH1.0 in an on-dyad conformation. ^33^ Missing residues and nucleotides were added using Modeller via the Chimera graphical user interface. ^65,66^ Linker histone coordinates from the 4QLC structure were used for GH1.0 on-dyad simulations, whereas for genGH1 on-dyad simulations the GH1.0 primary sequence was mutated to the genGH1 sequence (Figure 1). The completed on-dyad GH1 had an RMSD of 0.33 Å relative to the recently published crystallographic genGH1 structure (PDB 5NL0, resolution: 5.4 Å). ^34^ For simulations of the nucleosome, the linker histone was deleted.

Off-dyad binding models were based on a combination of manual placement, rigid docking, and flexible docking. First, the exit DNA was manually adjusted to allow space for the linker histone to be placed in an off-dyad binding mode. Rigid docking of genGH1 was then performed with the 12 Å cryo-EM map as a guide using the Colores module of Situs, ^37,67,68^ which was followed by flexible docking using internal coordinates normal mode analysis (iMOD). ^69^ To validate the linker histone placement, theoretical paramagnetic relaxation enhancement (PRE) intensity ratios were calculated and compared to experimental data (see Analyses Methods below). A model of off-dyad GH1.0 binding was constructed by superimposing and replacing genGH1 coordinates with GH1.0 coordinates.

### Molecular Dynamics Simulations

All systems were prepared with *tleap* from the AmberTools16 ^70^ software package. Each system was solvated in a TIP3P water box extending at least 10 Å from the solute. ^71,72^ Using Joung-Cheatham ions, ^73,74^ the solvent contained 150 mM NaCl, sodium cations to neutralize negative charges, and magnesium ions that replaced the manganese ions in the 1KX5 crystal structure. Only magnesium ions in the DNA grooves were included, whereas those located close to the the linker histones binding locations were excluded so as to not interfere with LH-DNA interactions. The AMBER14SB and BSC1 force fields were used for protein and DNA interactions. ^75,76^ All simulations were performed using NAMD version 2.12. ^77^ A cutoff distance of 10.0 Å with a switching function beginning at 8.0 Å was used for nonbonded interactions, and long range electrostatics were treated with particle mesh Ewald calculations. ^78^ For constant pressure calculations a modified NAMD version of the Nosé-Hoover barostat was used with a target pressure of 1.01325 bar while the Langevin thermostat was with a target temperature of 300K. ^79^

Systems were minimized twice for 5000 steps, first with a 10 kcal·mol^-1^·Å^-2^ harmonic restraint applied to the solute and then followed by no restraints. Using the Langevin thermostat, systems were then heated in the NVT ensemble from a temperature of 10K to 300K in 1K increments every 4 ps with a 10 kcal/mol kcal·mol^-1^·Å^-2^ solute restraint. This restraint was then reduced by 0.001 kcal/mol every 60 femtoseconds in the NPT ensemble. Equilibration runs with no restraints and a temperature of 300K were then performed. Simulations were conducted on seven systems: four chromatosomes (each containing genGH1 or GH1.0 in either the on- and or off-dyad binding mode), one nucleosome, and two isolated linker histones (genGH1 and GH1.0). Each simulation was run in triplicate for 250 ns using resources provided by the Extreme Science and Engineering Discovery Environment (XSEDE). ^80^

#### Umbrella Sampling

The ϕ_2_ reaction coordinate was divided into 41 windows spaced every 2 degrees, which covered a range of 40 to 120 degrees. Seed structures for each window were selected from steered MD runs where ϕ_2_ was adjusted at a rate of 5 deg/ns using a harmonic force constant of 0.1 kcal/mol from the closed-state in the 1HST structure. ^55^ The angle used here, along with ϕ_1_, is detailed in Figure S2 of Section S4 and Table S3 of Section S3. Windows were run for 20 ns each with a force constant of 0.1 kcal/mol/deg^2^, totaling 1.64 μs of simulation data, where the first 5ns of simulation was removed from analysis for equilibration. MD simulation parameters in NAMD were the same as described above for simulations of linker histones in solution, with the umbrella potential provided by the colvars module. ^81^ Trajectories were analyzed using the weighted histogram analysis method (WHAM) ^82^ with code from the Grossfield group. ^83^ For the Monte Carlo bootstrapping error analysis, 100 trials were run for each distribution and the statistical inefficiency was calculated for each window to calculate the number of statistically independent data points in each window. Convergence of each PMF can be found in Figure S3 of the SI.

### Analyses Methods

Unless otherwise stated, all analyses were performed using *cpptraj* with the first 50 ns of the trajectory excluded for equilibration. ^70^ Figures were created using Visual Molecular Dynamics. ^84^ Protein-DNA contacts were defined between the heavy atoms of residues within 4.0 Å of one another.

#### Estimated Binding Affinity

Binding affinities were estimated with an MM/GBSA analysis (molecular mechanics - generalized Born surface area) using the *MMPBSA.py* script from the AMBER16 software suite. ^85^ The three trajectory approach was implemented by using the trajectories from chromatosome simulations as the complex, the nucleosome simulations as the receptor, and simulations of linker histones in solution as the ligands. Explicit solvent trajectories were stripped of all solvent molecules while using trajectory frames every 4 ps. The implicit solvent model, *GBneck2* with the *mbondi3* radii parameters were used as they have been shown to have good agreement with more expensive Poisson–Boltzmann calculations for protein/nucleic-acid complexes. ^86^ The salt concentration was set to 150 mM.

#### Clustering

Binding modes of genGH1 and GH1.0 were compared by calculating RMSD values of the helical α-carbons in the linker histone with respect to the helical α-carbons in the core histones. The RMSD analysis was limited to the helical α-carbons to reduce noise from the loops and intrinsically disordered tails. Based on the RMSD results, a cutoff of 2.0 Å was chosen for the subsequent clustering analysis (Figure S4). Clustering was performed using the hierarchical agglomerative approach implemented in the *cluster* module of *cpptraj* from the the AmberTools16 software package. ^87^

#### Linker DNA Dynamics

The linker DNA in- and out-of nucleosomal plane motions were quantified to describe the linker DNA motions. To define the plane, the nucleosomal DNA was divided into four quadrants and the center of mass of the C1’ atoms within the two quadrants located distal from the linker DNA were used for two points, while the third point was defined as the C1’ center of mass of bases 83 and 250 which are located approximately on the dyad axis (see section S2 for details). The linker DNA vectors were defined as the C1’ center of mass of the base pairs at the origin of the linker DNA (bases 20-315 and 148-187) and terminal base pairs (bases 1-334 and 167-168), respectively. The α-angles were defined as in-plane and the β-angles were defined as out-of-plane motions of this vector. Positive α-angles were defined as inward motions towards the dyad axis while positive β-angles were defined as outward motions away from the nucleosomal-plane. For reference, the angles shown in Figure 6 are positive.

**Figure 6:**
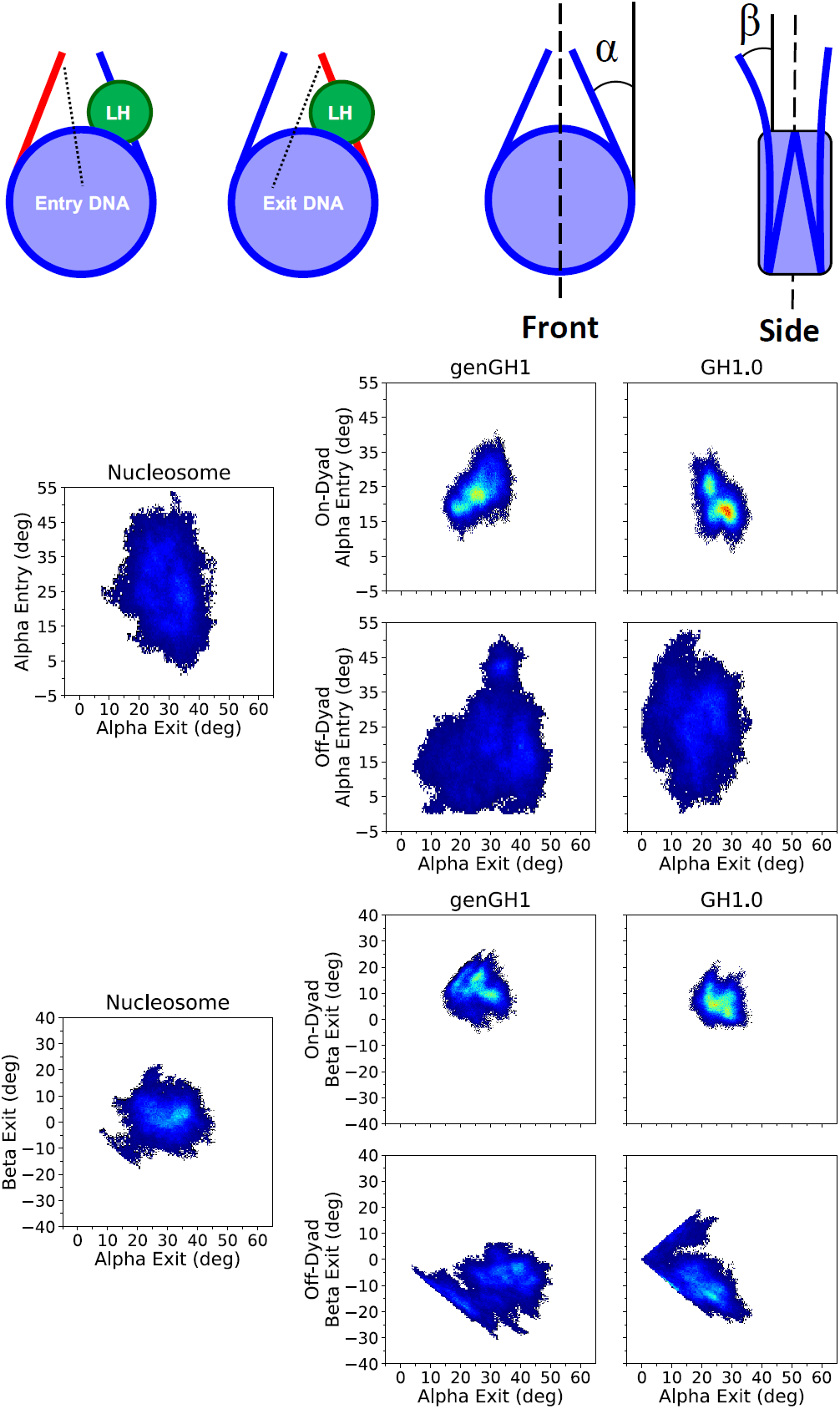
Density plots of linker DNA α and β angles. Entry and Exit DNA (from left to right) are defined in the top left graphic. The linker histone (LH) is shown in green, the linker DNA arm in red, while the rest of the nucleosome is blue. To the right are the α and β definitions (from left to right), which were inspired by Bednar *et al.* ^34^ The dyad axis is shown as a black dotted line. Each plot shows a 2D histogram of the entry linker DNA angles versus the exit linker DNA angles for both α (middle set of plots) and β (bottom set of plots) angles, with density ranges from dark blue (lowest) to red (highest). Each plot contains the linker DNA from the nucleosome core particle along with genGH1 and GH1.0 in the on- and off-dyad binding modes.

The normalized mutual information (NMI) between angles was calculated to determine the correlations between each pair of angles. ^88^ The NMI has the advantageous property that all values are in the range of 0 to 1 and are therefore easier to interpret than standard mutual information (MI). For detail discussion of the NMI see Supporting Information. The change in sampling of bound linker histones from the nucleosome were computed by the Kullback–Leibler divergence utilizing the nucleosome sampling distributions as the reference set: ^89^

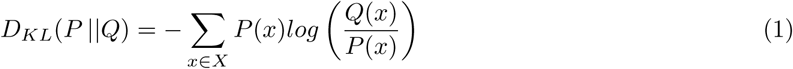

where *Q*(*x*) is the normalized reference distribution (nucleosomal linker DNA angles) and *P* (*x*) is the normalized data set (chromatosomal linker DNA angles).

#### Paramagnetic relaxation enhancement (PRE) intensity ratios

Theoretical paramagnetic relaxation enhancement (PRE) cross-peak intensity ratios were estimated based on distances between experimentally labelled core histone (H2A T119 and H3 K37) and linker histone methyl-terminated residues. ^36,38^ Since our simulations did not include the MTSL probe, wild type residues were used for distance calculations, similar to Piana *et al.* ^90^ For the K37 probe, distances were measured from the lysine terminal nitrogen to the terminal methyl carbon atom of each respective residue, while the T119 distance was measured from the threonine terminal methyl carbon. Note, that each of these neutral residues have two distinct methyl groups and were referred to as *methyl-a* and *methyl-b*. Using these distances, the predicted intensity ratios where calculated using known equations and the average of the *methyl-a* and *methyl-b* values were used to compared to experimental values.^36,38,62^

The relationship between between the paramagnetic relaxation cross-peak intensity ratios and the interatomic distances is given by:

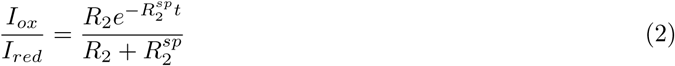

where, *R*_2_ is the intrinsic relaxation rate (inverse of the transverse time constant (*T*_2_)), *t* is the total evolution time of the transverse proton magnetization, *R*_*sp*_ is the contribution to the relaxation caused by the paramagnetic probe, and *I*_*ox*_ and *I*_*red*_ are the peak intensities for the oxidized and reduced states, respectively. This last variable is what connects the cross peak equation to the interatomic distances, *r*. Distances are defined by the following: ^91,92^

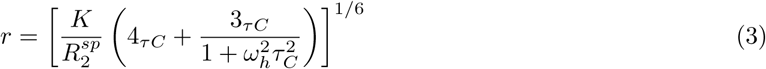

where, *K* is a constant that describes the spin properties of the MTSL label (1.23×10^−32^ cm^6^ sec^−2^), ω_*h*_ is the Larmour frequency, and τ_*C*_ is the apparent correlation time which is estimated from the molecular weight of the protein. All values are either known values from the experiment or constants.^33,36,38^ Zhou *et al.* further condensed equations (2) and (3) into a single equation:^33,36,38^

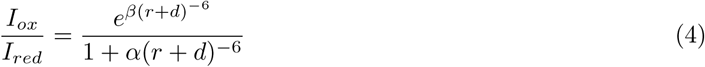

where, α= 4.5×10^8^, β= 3.4×10^7^, and *d* = 9.0 Å, a correction factor based on the experimental calibration curve.^33,36,38^ This is the equation we used to convert relevant distances in the structures and simulations to PRE cross peak intensity ratios.

## Results

### Unbound Linker Histones Favor the Closed-State

To quantify the dynamics of linker histones in solution, we measured two angles over our unbound genGH1 and GH1.0 simulations, ϕ_1_ and ϕ_2_, which were inspired by previous work on GH1.0 by Öztürk *et al.* (Figure 2, left side). ^53^ In that work, GH1.0 was shown to preferentially adopt the closed conformation in solution, although the open-state was sampled in accelerated MD simulations. Similarly, our unbound genGH1 and GH1.0 simulations predominantly sampled closed β-loop states. Both ϕ-angles are a measurement of the angle between the α3-helix and the β-loop with respect to either the α-helix (ϕ_1_) or the β-sheet (ϕ_2_). ϕ_1_ displayed a wider distribution than ϕ_2_ and often included values associated with both the open- and closed-states. For example, in genGH1 simulations there was a spread of ϕ_1_ from 54.6° to 125.4°, which encompasses both the crystolgraphic closed and open values of 116.2° and 96.9°, respectively. However, ϕ_2_ angles were more indicative of β-loop dynamics with average values of 52.6° ± 5.2° and 60.2° ± 8.9° for genGH1 and GH1.0, both of which correspond to closed-states. Therefore, ϕ_2_ appears to be a stronger metric of β-loop structures, whereas ϕ_1_ is subject to increased mobility throughout the β-sheet.

**Figure 2:**
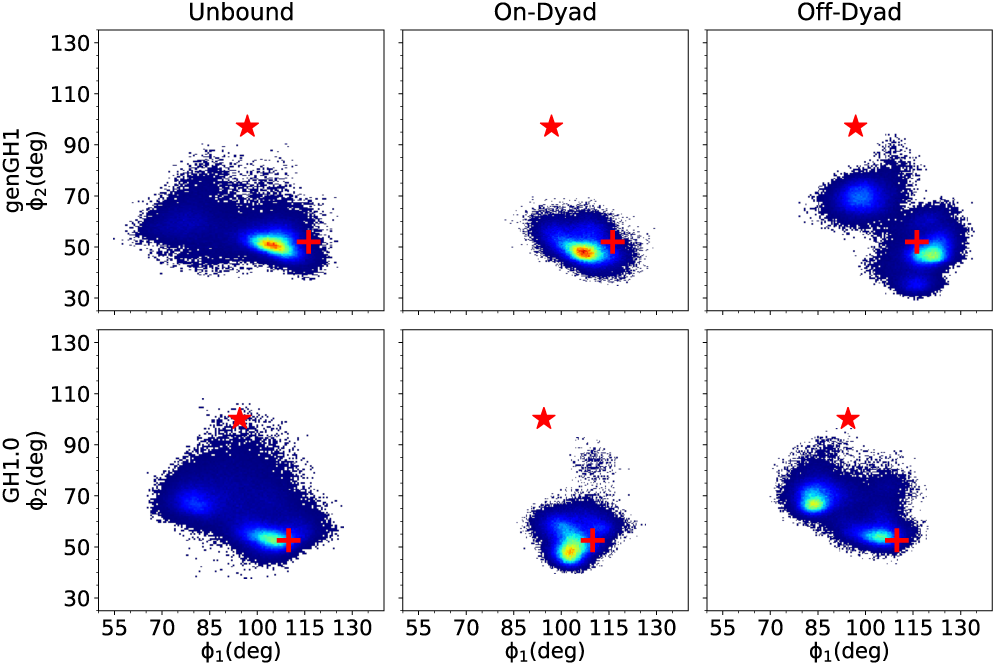
Flexibility of the β-loop using ϕ-angles inspired by a previous study. ^53^ Shown are 2D-histograms of ϕ_2_ versus ϕ_1_ angles (deg) in the unbound (left), on-dyad (middle), and off-dyad (right) states. Density ranges from blue (lower) to red (higher). The red cross corresponds to the initial values of the closed-state linker histone, while the red star corresponds to the open-state values.

To quantify the dynamics of the β-loop, umbrella sampling simulations were performed to determine the relative stability of the open-state in solution. Based on the results reported above, we determined that ϕ_2_ was a more direct metric of the β-loop dynamics than ϕ_1_. Consequently, here we computed potentials of mean force (PMF) along the one-dimensional ϕ_2_ coordinate space for both genGH1 and GH1.0 (Figure 3).The PMFs shows that both genGH1 and GH1.0 contain a single broad free-energy well spanning about 25° with minima at ∼50° and ∼62°. Based on the crystallographic structure, these states are consistent with closed β-loop. Free energies corresponding to the open β-loop (∼100°) suggest that this state is sparsely populated in solution for GH1.0, with a free energy of 3.27±0.48 kcal/mol, and virtually nonexistent for genGH1 which has an open-state free energy of 6.97±0.56 kcal/mol. For both systems, angles below 45° led to unphysical steric clashes, which results in increased free energies for these angles.

**Figure 3:**
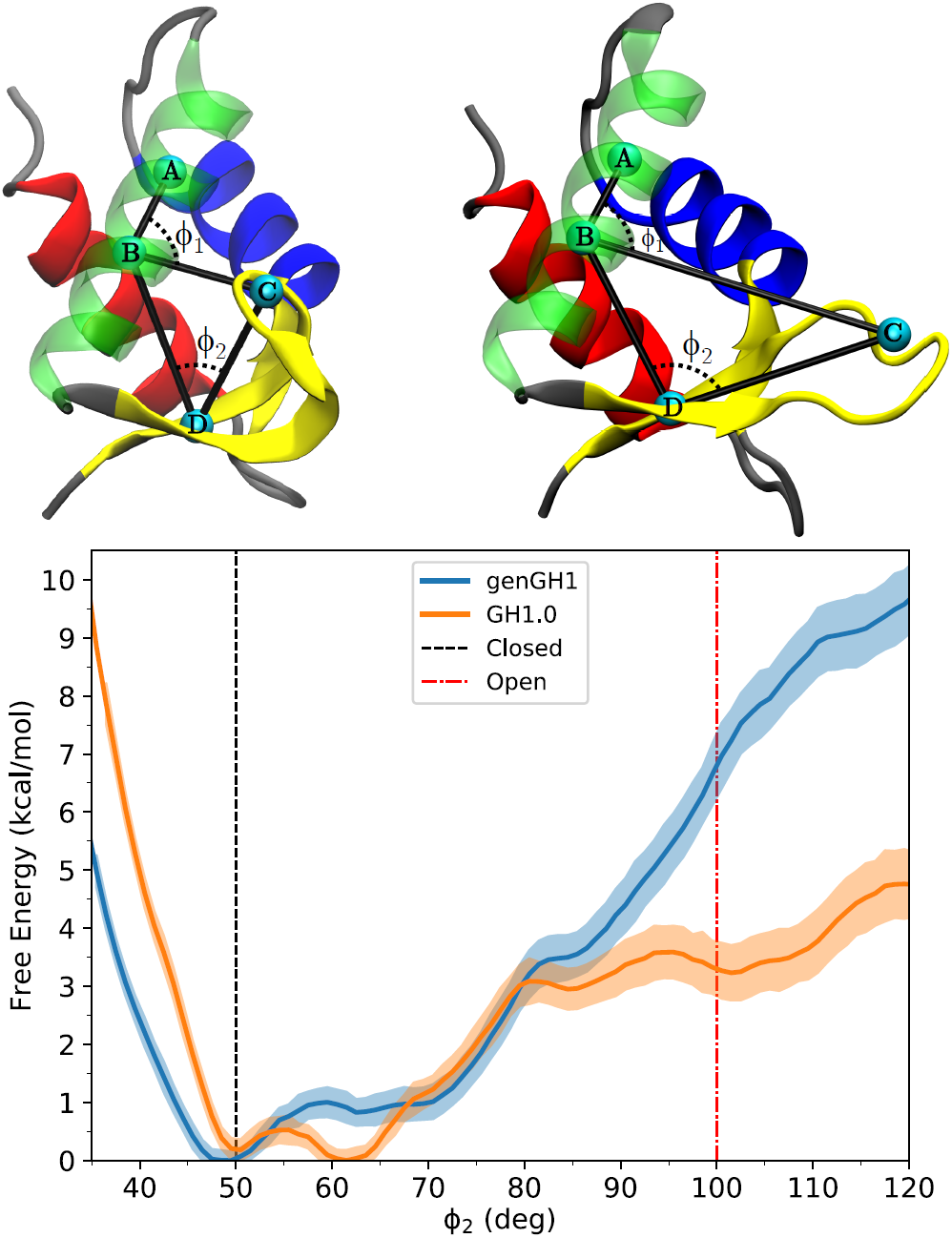
Potential of mean force from umbrella sampling simulations showing the relative free-energy landscapes between genGH1 and GH1.0 along the ϕ_2_ coordinate space. The red dotted-line corresponds to the closed-state angle, ϕ_2_ = ∼50.0°, while the green dotted-line corresponds to the open-state angle, ϕ_2_ = ∼100.0°, of the crystalographic closed-state from 1HST.

### Bound Linker Histone Dynamics

#### Off-Dyad Linker Histones Sampled Multiple States in the DNA Binding Pocket

There is no reported high-resolution crystal structure for linker histones bound in an off-dyad state. Therefore, a model for off-dyad genGH1 was constructed based on both the low-resolution cryo-EM map of a poly-nucleosomal array by Song *et al.* and NMR paramagnetic relaxation enhancements (PRE) experiments by Zhou *et al.* (see Methods). ^36–38^ Due to the availability of of the higher resolution PRE data, we used the same genH1 as Zhou *et al.* instead of the H1.4 represented in the array cryo-EM map and the H1.5 from the 5NL0 structure, and we assumed that off-dyad states are similar between linker histones, as has been observed for on-dyad states. The GH1.0 off-dyad model was constructed by mutating the genGH1 model to the appropriate primary sequence. This model was validated by comparing to PRE experiments conducted by Zhou *et al.* To make a direct comparison to the experimental data, we estimated the observed PRE intensity ratios (I_*ox*_/I_*red*_) using distances between the core histone labeled probed residues H2A T119 and H3 K37 and genGH1 methyl-terminated residues (Figure S5). In general, we observed suitable agreement for both sites, with mean signed errors of 0.02 and 0.04 for the T119 and K37 probe sites, respectively. Comparing our off-dyad model of the chromatosome to that described by Zhou *et al.*, the linker histones exhibit a slight rotational variation in the DNA pocket. ^38^ We attribute this difference largely to our use of data released after the work cited above, such as the cryo-EM map of the nucleosome array that was used to adjust the linker DNA arms. ^37^ Furthermore, our model exhibits more stabilizing LH-DNA contacts (see Figures S6 - S9) leading to an overall lower energy structure. Based on these results, we found the off-dyad structures of genGH1 and GH1.0 sufficient to begin simulations. For reference, we have provided the contacts of genGH1 and GH1.0 with the DNA prior to simulations (Figures S8 - S9).

For both genGH1 and GH1.0, there was considerably more sampling of the linker histone position in the off-dyad location relative to on-dyad (Figure S10). Specifically, the linker histone was localized within the DNA minor groove where it rocked in and out of the nucleosomal plane, although in our simulation it did not slide along the DNA between the on- and off-dyad states. This is quantified by the increased root-mean-square deviation (RMSD) values in off-dyad simulations (Figure S4). This contrast in RMSD values between binding modes is reflected in a clustering analysis which demonstrates that off-dyad systems have more clusters that are less populated than on-dyad systems (Figure 4). More specifically, when clusters were separated by 2 Å from one another, six clusters were found for GH1.0 in an on-dyad binding mode, whereas 11 were found for off-dyad GH1.0. Similarly, six and eight cluster were found for genGH1 on-dyad and off-dyad. However, some of these clusters had a low population, and when the clusters representing only the top 90% of frames were analyzed this resulted in three clusters for each on-dyad and five clusters for each off-dyad linker histone (Figure 4). These suggest that off-dyad linker histones are more fluid in the DNA pocket. In contrast, the increased number of linker histone-DNA contacts in the on-dyad pocket (discussed below) leads to a more confined and rigid complex, hence decreased sampling.

**Figure 4:**
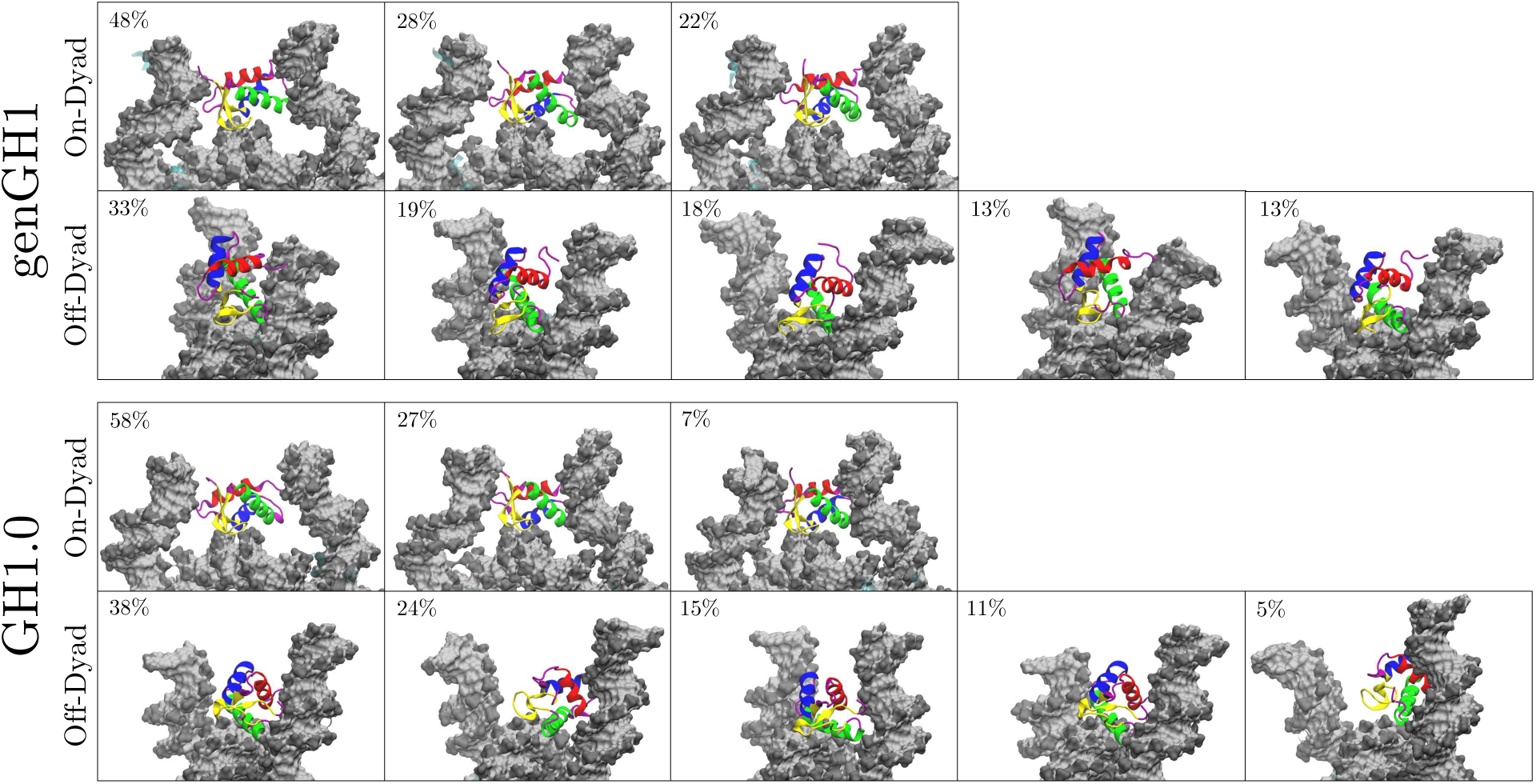
Representative snapshots of clusters for linker histones genGH1 and GH1.0 in both the on- and off-dyad binding modes. Shown are clusters totalling at least the top 90% of frames analyzed. The respective percentage of each cluster is provided in the top left-hand corner of each snapshot.

This increased sampling in off-dyad systems contributes to increased uncertainty in the genGH1 observed time-averaged PRE ratios (Figure S11 and S12 for GH1.0). These values had a higher discrepancy to experiments with mean errors of 0.11 and 0.16, although for most residues the experimental values were within the 80% confidence interval of the simulation derived ratios. Some of these differences are likely due to localized fluctuations of the linker histones that occur throughout the simulations and the fact that distance changes on the order of 2-3 Å in methyl/probe distances can result in difference on the order of 0.10-0.15 in the PRE intensity ratio.

In all bound linker histones simulations the β-loop remained in the closed conformation. This is exemplified by the ϕ_2_ angle distributions, which sampled states that had a free energy in solution below 1 kcal/mol with values of 49.7° ± 4.2° and 55.0° ± 11.5° for genGH1 in on- and off-dyad states, and 52.4° ± 5.6° and 63.7° ± 7.9° for GH1.0 on- and off-dyad systems (Figure 2). As previously noted, ϕ_1_ is a relatively poor metric to distinguish between the open- and closed β-loop states.

#### On-Dyad Binding Restricts DNA Motions

Linker histones interact with both the nucleosomal and linker DNA, which has a direct effect on their motions within the DNA binding pocket. Above, we have emphasized the importance of this interaction by showing how the linker histone binding pose affects experimental results. To further probe these dynamics, we plotted the in- and out-of-nucleosomal-plane motions of both linker DNA arms (Figure 5 and S13). Generally, in on-dyad binding modes the interactions of the linker histone with both linker DNA arms restricted both of their motions compared to the nucleosome. However, when the linker histone was bound off-dyad, the entry-linker DNA sampling was similar to the nucleosome, whereas the exit-DNA was shifted out of the nucleosomal plane. Due to the asymmetric nature of the Widom 601 DNA sequence, DNA motions were not symmetric in the nucleosome simulations between the entry and exit DNA segments. Beyond Figure 5, we further quantified the in- and out-of-plane linker DNA motions which are termed here as the α and β angles, respectively, for the entry and exit DNA, as inspired by Bednar *et al.* ^34^ (see Figures 6 and S1 for definitions). The α angles correspond largely to DNA breathing motions, and over all simulations ranged from 0.0° to 54.0° with an average value of 25.0°. The lack of negative α angles indicates that, on the timescales sampled here, no significant opening of the DNA was observed. Out-of-plane linker DNA fluctuations, described by β angles, ranged from −27.4° to 32.1° in all simulations, similar in scope to the α angles. These angles are complimented by DNA-end-to-end distances in Figure S14.

**Figure 5:**
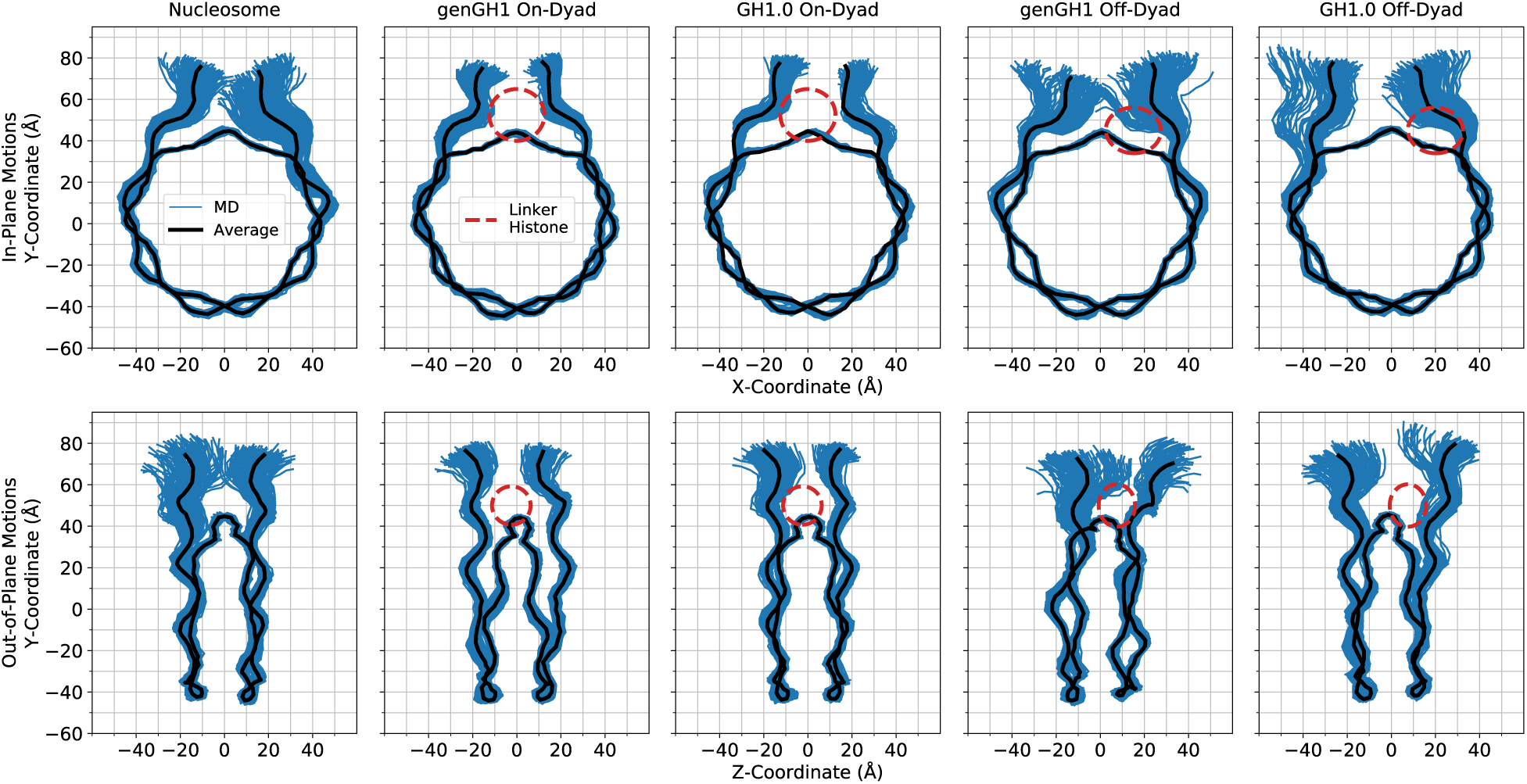
In-plane (top) and out-of-plane (bottom) DNA motions sampled by the genGH1, and GH1.0 in the on- and off-dyad binding modes along with the nucleosome. Shown in blue are configurations sampled throughout the MD simulation (150 representative frames) while the average configuration is shown in black. For reference, the approximate position of the linker histone is shown as a dashed-line red ellipse. Figures inspired by work from Shaytan *et al*. ^93^

Linker histone binding had a significant effect on both the equilibrium distributions of these motions as well as their correlations to one another. In addition to the 2D-histograms of the α and β angles (Figures 6 and S15 - S16), their correlations were analyzed by computing the normalized mutual information (NMI) for each angle pair (Table 1), and the changes induced by linker histone binding were quantified with the Kullback-Leibler (KL) divergence for each probability distribution relative to the nucleosome (Tables 2 & S2). Of particular note are the α-entry/α-exit angle distributions which highlight the breathing motions of both DNA ends (Figure 6, middle). In this space, the nucleosome samples a wide range of angles in both the entry and exit DNA with little correlation between the two as shown by the NMI value of 0.04. In off-dyad binding there is a similar range of motions for genGH1, which has a KL divergence of only 1.62 relative to the nucleosome, while GH1.0 has a similar range of sampled states with a shifted mean, resulting in a higher KL value of 3.95. For both cases the correlations between the α angles are relatively low, with a modest increase in the NMI values for genGH1 and a slight decrease for GH1.0. In contrast, on-dyad binding shows a significant reduction in the α-entry/α-exit conformational space, as the range of motion of both the entry and exit DNAs are substantially restricted regardless of the linker histone variant. The KL divergences for these states are high, 4.35 and 5.82 for genGH1 and GH1.0, and the correlations for each state are approximately three times higher than for the nucleosome.

**Table 1:**
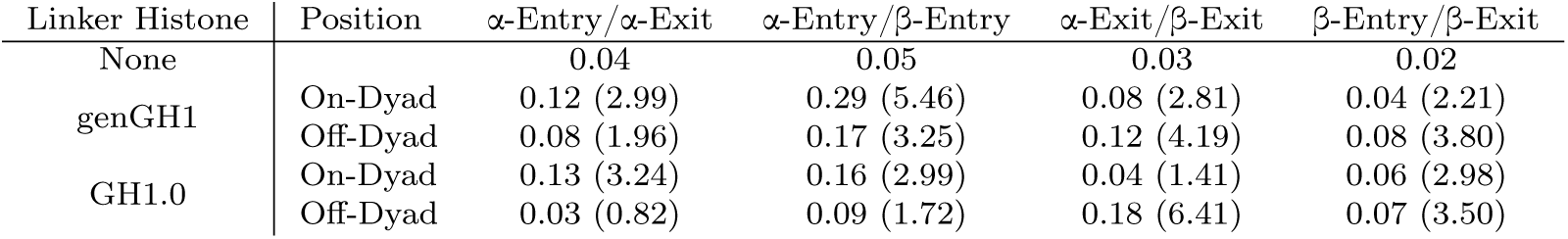
Normalized mutual information values for DNA motions. Values in parenthesis are the increase in the mutual information for that measurement over the canonical nucleosome.

**Table 2:** Kullback-Leibler divergence values for two dimensional probability distributions of DNA. Cells are colored on a scale from blue (lower) to white to red (higher).

| Linker Histone | Position | α-Entry/α-Exit | α-Entry/β-Entry | α-Exit/β-Exit | β-Entry/β-Exit |
| --- | --- | --- | --- | --- | --- |
| genGH1 | On-Dyad | 4.35 | 5.55 | 4.56 | 5.52 |
|  | Off-Dyad | 1.62 | 1.75 | 6.33 | 6.11 |
| GH1.0 | On-Dyad | 5.82 | 5.12 | 3.01 | 1.91 |
|  | Off-Dyad | 3.95 | 1.58 | 9.48 | 3.36 |

Another distribution of interest is the α-exit/β-exit phase space, which describes the in- and out-of-plane motions of the exit DNA (Figure 6, bottom). In the canonical nucleosome the average α-exit and β-exit angles are 30.5° and −2.7° respectively, indicating that this DNA arm fluctuates about states that are slightly pointing inward when viewed from the side. These motions are largely uncorrelated, with an NMI value of 0.03. Binding in the off-dyad location has a dramatic effect, forcing the DNA outward and shifting the β-exit angles to fluctuate around 9.5° and 7.1° for the genGH1 and GH1.0 systems. This alters the sampling distributions, with KL values of these angles of 6.33 and 9.48 for genGH1 and GH1.0, and increasing the correlations between the motions by four to six fold. In contrast, on-dyad binding decreases the average β-exit value to −12.1° and −6.1° for genGH1 and GH1.0 as this binding mode pulls the DNA inward from the side view. This results in more modest changes to the α-exit/β-exit phase space and lower correlation increases over the nucleosome.

In contrast to the exit DNA, the entry DNA is largely unaffected by off-dyad binding with KL values below 1.8 relative to the nucleosome. On-dyad binding creates a larger perturbation, with KL values above 5.1 as ranges of motion of both angles are restricted relative to the nucleosome. The average β-exit angles have a slight increase from −5.1° in the nucleosome to 1.8° and 0.6° in genGH1 and GH1.0, which are both significantly greater than the values in the exit DNA. Together, this creates an asymmetry in the profile view of DNA structures, as illustrated in Figure S15.

### Energetic Contributions

#### On-Dyad Binding is Enthalpically Favored

Although both linker histones have similar physical effects on the nucleosome when bound in on- or off-dyad locations, the difference between their binding energetics determines their *in vitro* binding mode preference. To estimate binding affinities, we have used an MM/GBSA analysis to decompose the overall binding affinity into the energetic components that drive this interactions. ^94^ Regardless of the variant, on-dyad binding was found to be significantly more enthalpically favorable than off-dyad binding (Table 3). GH1.0 favored the on-dyad state over off-dyad by −163.3 ± 36.3 kcal/mol, while genGH1 only favored on-dyad by −89.9 ± kcal/mol. This difference in ΔΔE_total_ values was largely driven by Van der Waals (VdW) interaction energies, with GH1.0 favoring the on-dyad state by −91.0 ± 24.5 kcal/mol and genGH1 favoring off-dyad state by 32.1 ± 25.6 kcal/mol. In contrast, genGH1 favors the on-dyad binding mode more than GH1.0 in the electrostatic interaction energies. However, a large overlap in the errors between variants suggests that both systems have a relatively similar electrostatic on-dyad binding preference. We emphasize that MM/GBSA approach includes a number of approximations and do not include important thermodynamic quantities such as conformational entropy or explicit solvent thermodynamics, therefore these values should be taken as qualitative estimates of binding affinities. ^95,96^

**Table 3:** Energy differences (kcal/mol) between on and off-dyad binding states as estimated by MM/GBSA analysis. A negative value indicates more favorable binding in the on-dyad state.

| Linker Histone | Binding Mode | $\Delta E_{\text{total}}$ | $\Delta E_{\text{internal}}$ | $\Delta E_{\text{elec}}$ | $\Delta E_{\text{VdW}}$ | $\Delta\Delta E_{\text{total}}$ | $\Delta\Delta E_{\text{internal}}$ | $\Delta\Delta E_{\text{elec}}$ | $\Delta\Delta E_{\text{VdW}}$ |
| --- | --- | --- | --- | --- | --- | --- | --- | --- | --- |
| genGH1 | On-Dyad | $-174.6 \pm 37.0$ | $-18.1 \pm 32.5$ | $-59.5 \pm 27.2$ | $-97.0 \pm 23.9$ | $-89.9 \pm 39.0$ | $-29.1 \pm 33.3$ | $-92.9 \pm 29.4$ | $32.1 \pm 25.6$ |
| | Off-Dyad | $-84.5 \pm 39.9$ | $11.0 \pm 33.1$ | $33.4 \pm 28.3$ | $-128.9 \pm 23.6$ | | | | |
| GH1.0 | On-Dyad | $-221.6 \pm 37.8$ | $-13.4 \pm 32.2$ | $-61.9 \pm 24.7$ | $-146.4 \pm 24.4$ | $-163.3 \pm 36.3$ | $-3.3 \pm 33.0$ | $-69.0 \pm 22.8$ | $-91.0 \pm 24.5$ |
| | Off-Dyad | $-58.2 \pm 36.5$ | $-10.1 \pm 33.1$ | $7.1 \pm 24.2$ | $-55.3 \pm 22.0$ | | | | |

#### Van der Waals Interactions Drive Binding Mode Selectivity

To further investigate the thermodynamic driving forces between linker histone variants and binding locations, we calculated the energetic strain between each binding species in their complexed and isolated states, as defined by:

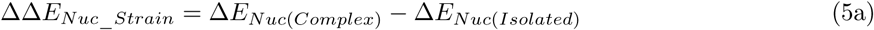

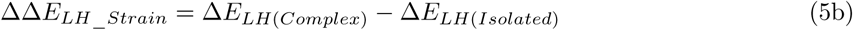

The results in Table 4 suggest that a combination of electrostatic and VdW interactions from both the linker histones and nucleosomes drive the system conformations. The ΔΔE_tot_ of genGH1 for on-dyad (17.9 ± 5.2 kcal/mol) and off-dyad (28.2 ± 5.6 kcal/mol) is ∼2.5-fold greater than GH1.0 on-dyad (7.4 ± 5.0 kcal/mol) and off-dyad (10.6 ± 5.3 kcal/mol). These variations are largely defined by differences in the VdW interactions where the ΔΔE_VdW_ of GH1.0 is 14.9 kcal/mol and 17.2 kcal/mol more favorable than genGH1 for on- and off-dyad systems, respectively. Additionally, it is worth noting that electrostatic interactions (ΔΔE_ele_) for both linker histones actually favor the off-dyad complex by 11.6 kcal/mol and 19.9 kcal/mol for genGH1 and GH1.0, respectively.

**Table 4:** Energetic strain (kcal/mol) between binding species while each are in complex and in isolation as estimated by MM/GBSA analysis. A negative value (-) indicates a favorability of the binding species in complex, as oppose to an isolated state, positive (+). Full binding energies for each state are given in Table S1.

| Linker Histone | Binding Mode | Binding Species | $\Delta\Delta E_{tot}$ | $\Delta\Delta E_{int}$ | $\Delta\Delta E_{ele}$ | $\Delta\Delta E_{VdW}$ |
| --- | --- | --- | --- | --- | --- | --- |
| genGH1 | On-Dyad | Nuc | $-74.8 \pm 25.0$ | $-15.9 \pm 23.0$ | $-81.1 \pm 20.0$ | $22.2 \pm 17.7$ |
| | | LH | $17.9 \pm 5.2$ | $-2.3 \pm 5.0$ | $13.3 \pm 2.8$ | $6.8 \pm 3.0$ |
| | Off-Dyad | Nuc | $-40.3 \pm 29.2$ | $8.4 \pm 23.6$ | $-29.1 \pm 20.7$ | $-19.6 \pm 16.4$ |
| | | LH | $28.2 \pm 5.6$ | $2.5 \pm 4.9$ | $1.7 \pm 2.9$ | $24.0 \pm 3.5$ |
| GH1.0 | On-Dyad | Nuc | $-95.8 \pm 26.0$ | $-12.0 \pm 22.8$ | $-74.6 \pm 16.5$ | $-9.2 \pm 17.8$ |
| | | LH | $7.4 \pm 5.0$ | $-1.4 \pm 4.9$ | $17.0 \pm 3.1$ | $-8.1 \pm 2.7$ |
| | Off-Dyad | Nuc | $5.7 \pm 24.8$ | $-16.8 \pm 24.0$ | $-44.9 \pm 15.3$ | $67.5 \pm 15.5$ |
| | | LH | $10.6 \pm 5.3$ | $6.6 \pm 5.0$ | $-2.9 \pm 2.8$ | $6.8 \pm 2.9$ |

Off-dyad nucleosomes showed an increased stability when bound to genGH1 over GH1.0 with a ΔΔE_tot_ of −40.3 ± 29.2 kcal/mol and 5.7 ± 24.8 kcal/mol for genGH1 and GH1.0 systems, respectively. This contrast in ΔΔE_tot_ can be also be attributed to the VdW energies with ΔΔE_VdW_ values of −19.6 ± 16.4 kcal/mol and 67.5 ± 15.5 kcal/mol for genGH1 and GH1.0 systems, respectively.

Contacts between linker histones and the DNA (Figure S17) show that most on-dyad interactions come from the β-sheet (Figure 7), with 66.8 ± 12.9 and 71.2 ± 11.8 contacts for genGH1 and GH1.0. However, the contrast between variants becomes more evident in the off-dyad binding mode with 30.2 ± 14.8 and 2.2 ± 2.6 β-sheet-DNA contacts for genGH1 and GH1.0, respectively. A similar relationship is also observed in the off-dyad α3 helix and N’-tail. The genGH1 α3 helix in the off-dyad binding mode has 16.4 ± 10.8 more contacts than GH1.0 while the N’-tail has 9.1 ± 18.0 more contacts. Combined, these additional contacts between genGH1 and DNA correlate with its VdW-driven preference for off-dyad binding over GH1.0. Taken together, these results suggest that while on-dyad binding is energetically preferred in both variants, genGH1 has a higher propensity for sampling the off-dyad state than GH1.0, which is largely due to the difference in contacts between the β-sheet and the DNA.

**Figure 7:**
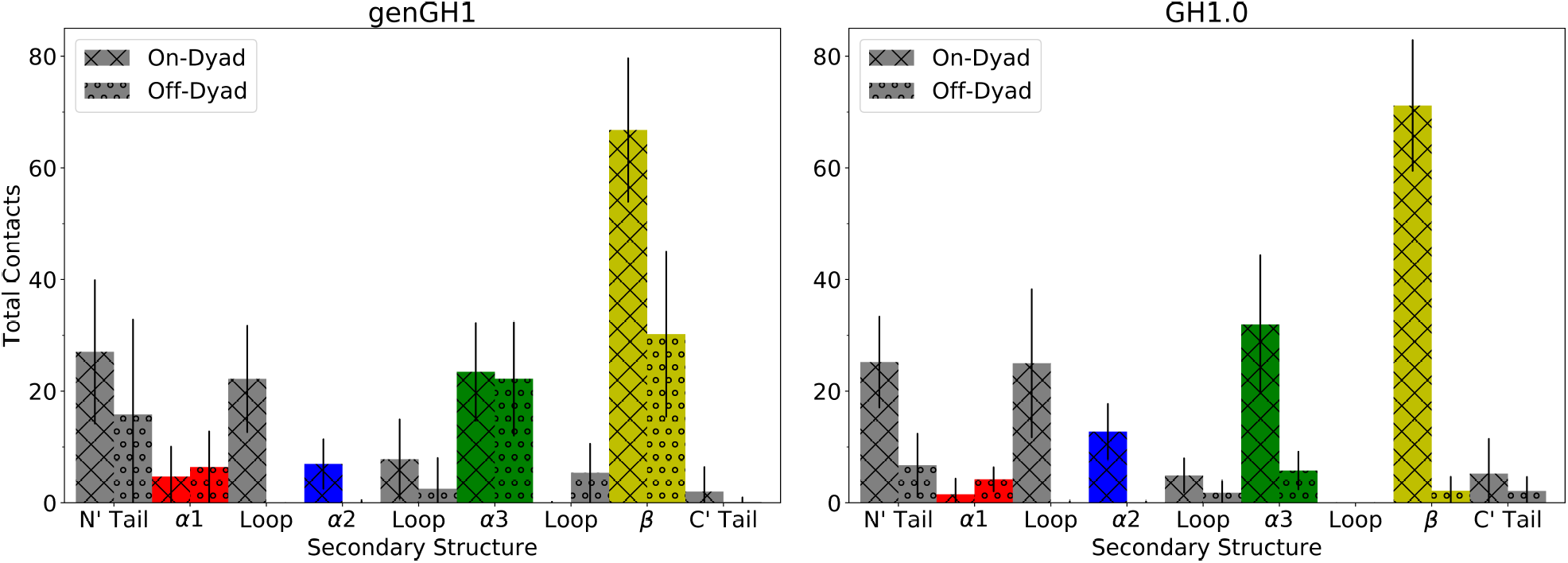
Total contacts of genGH1 and GH1.0 secondary structures with DNA in both the on- and off-dyad binding modes. The secondary structure is broken down into the following: alpha helix α1 (red), alpha helix α2 (blue), alpha helix α3 (green), β-sheet (yellow), N-terminal tail (grey), C-terminal tail (grey), and three loops (grey).

## Discussion

Here, we have used a combination of conventional and free energy molecular dynamics simulations to probe the effects of linker histone binding on chromatosome structures and dynamics. Umbrella sampling PMFs show that for both genGH1 and GH1.0 the closed-state is thermodynamically favorable, which is consistent with the GH1.0 accelerated MD results of Öztürk *et al.* ^53^ This suggests that the open-state in the 1HST crystal structure is stabilized by crystal packing forces, as exemplified by the fact that the β-loop is inserted into a hydrophobic pocket in the neighboring unit. These results are in general agreement with crystal structures of linker histones bound in the on-dyad state which have the β-loop in the closed-state.^33,34,46^

Given that there is no high resolution crystal structure of off-dyad binding, the precise binding structure and orientation of linker histones in this pocket remains inconclusive.^36,38^ We therefore constructed a model of the off-dyad state based on manual placement and docking into the 30-nm cryo-EM structure by Song *et al.* which we found had generally good agreement with the PRE data from Zhou *et al.* ^36,37^ Based on our umbrella sampling simulations we used the closed-states of the β-loops in these structures. As highlighted by the clustering results in Figure 4, our simulations show that both linker histones are significantly more fluid in the off-dyad DNA binding pocket relative to on-dyad. This is in line with the results of Brownian dynamics docking studies from the Wade group in which they found that genGH1, GH1.0, and assorted mutants can bind in a variety of sequence dependent orientations. ^54^ Furthermore, these series of simulation results would suggest transitions between on- and off-dyad states might be facilitated by multiple stable binding orientations along the DNA and encouraged by additional linker histone conformational freedom in the binding pocket.

One of the central mechanisms by which linker histones inhibit transcription and promote the compaction of chromatin fibers is by altering linker DNA dynamics. Our simulations have shown that one of the primary differences in on- and off-dyad binding is that on-dyad binding drastically restricts both the entry and exit DNA segments, whereas off-dyad binding has a distinct influence on the exit DNA dynamics with little change to the entry DNA. The latter is in line with previous Brownian and molecular dynamics simulations from the Wade group which have shown that GH1.0 modifies linker DNA motions in off-dyad binding modes. ^53,56^ Furthermore, our results indicate that these effects are largely independent of linker histone variant type. These differences in DNA dynamics have broad implications for greater chromatin structures. For example, Mishra and Hayes have highlighted how the stoichiometric binding of H1 to nucleosome arrays can severely limit linker DNA accessibility to trans-acting factors. ^31^ Meanwhile, cryo-EM structures of polynucleosome arrays have revealed linker histones in both on- and off-dyad locations, with distinct effects on the greater structure of chromatin arrays. ^37,42^ Furthermore, Perišić *et al.* recently showed with a highly coarse-grained model that linker histone binding position influences their tail positions, which directly impacts greater chromatin structures, with off-dyad linker histones creating more condensed chromatin fibers. ^41^ Finally, numerous studies have shown chromatin fiber pliability to be highly dependent on the linker DNA length and, by extension, the nucleosome repeat length. ^43–45^ In light of these results, it is likely that *in vitro* on-dyad mono-chromatosomes are a result of additional linker histone DNA contacts which may not be as prevalent in condensed nucleosome arrays due to limited linker DNA conformational freedom. ^34^

Given that genGH1 and GH1.0 have similar effects on the structure and dynamics of chromatin when bound in on- and off-dyad locations, what is the role of various linker histone variants and modifications *in vitro*? Results from our energetic and contact analyses show that while they have similar structures, the genGH1 and GH1.0 variants have drastically different energetic preferences for the on- and off-dyad states. Indeed, while both have an enthalpic preference for on-dyad binding, the preference is significantly reduced for genGH1. To further quantify the thermodynamics of binding would require explicit calculations of entropic contributions differences in each system. Unfortunately, entropic calculations on systems of this size are notoriously difficult to converge. ^97^ Therefore, we have extrapolated their influences based on changes in the linker DNA and linker histone dynamics. The results suggest that while the enthalpic reward of on-dyad binding in the GH1.0 chromatosome is enough to overcome the entropic penalty from the drastic reduction in linker DNA and linker histone dynamics, that in the case of genGH1 there is not enough of an enthalpic difference between on- and off-dyad states to compensate for these entropic losses, which is why genH1 has typically been observed in the off-dyad binding mode *in vitro*. This is in line with experimental results that consistently show GH1.0 in the on-dyad state, but that have provided evidence for genGH1 in both the on- and off-dyad states.^34,37,46^

In an excellent example of the finely-tuned nature of the on- and off-dyad binding thermodynamics, Zhou *et. al* used PRE NMR experiments to show that mutation of five GH1.0 residues can shift its binding equilibrium from on- to off-dyad locations. ^36^ We performed additional simulations of this GH1.0 pentamutant, and observed that dynamics in on- and off-dyad states are similar to those for the GH1.0 and gen1.0 systems examined above (Section S5: S5.1 - S54; Figures S19 - S23). However, these mutations did have a significant effect on the binding energies. Specifically, they lowered the favorability of the on-dyad state, while at the same time had little effect on the off-dyad binding energies (Table S4). This change in energy brings the on-dyad binding energy to within error of the off-dyad state.Taken with the increased dynamics in off-dyad states, likely resulting in an entropic penalty of on-dyad binding, this suggests the binding free energy favors the off-dyad state, as observed by Zhou *et al.*, and demonstrates how even relatively minor changes to linker histone structures can have significant influences on their binding thermodynamics.

More specifically, we observe that the change in energetic preference between the two states is driven by increased β-sheet, α3-helix, and N-terminal tail contacts in genGH1 off-dyad systems. Surprisingly, these changes in contacts contributes mostly to differences in the Van der Waals interactions, highlighting their often overlooked influence in protein-DNA binding thermodynamics. Although we emphasize that electrostatics are vital in the binding process, our results show them to be relatively consistent between the on- and off-dyad binding modes, independent of linker histone variant. In the off-dyad binding orientation of the linker histone, the mostly conserved β-sheet is more solvent exposed. Given that each linker histone has the same initial placement, the loss of contacts from the on-dyad state to the off-dyad should be similar, specifically between the β-sheet and DNA. However, the linker histones facets that remain in contact with the DNA exhibit a less conserved sequence which would express a greater difference in VdW energies due to sidechain variations. We further investigated this by examining the propensity of linker histone residues to be in contact with particular parts of each nucleotides, such as the backbone, sugar, and base. Generally, we found that contacts between the DNA backbone and linker histone residues were more dominant, except in the case of off-dyad genGH1 (Figure S18). In this system, residues in the β-sheet are consistently in contact with the hydrophobic bases of each nucleotide, overshadowing contacts with the backbone. Together, these results suggest that chromatosome systems are driven by a subtle equilibrium wherein multiple binding states may simultaneously be populated in solution. *In vitro*, there are likely transitions between the on- and off-dyad binding states, with linker histones diffusing along the DNA while guiding chromatin fiber flexibility. While the on-dyad enthalpic reward seen in the GH1.0 chromatosome may be strong enough to overcome the entropic penalty observed in the linker DNA and linker histone, this is not the case for genGH1 which is likely why it has more often been observed in the off-dyad binding mode.

The initial structures used for MD simulations come from a combination of X-ray diffraction (XRD) and cryo-EM structures. In general, cryo-EM-based structures are often more representative of *in vitro* systems than X-ray diffraction structures. Despite this, the DNA and linker histone of 4QLC^33^ (XRD) are very similar to the cryo-EM 5NL0^34^ structures. Moreover, the core histones of our models come from 1KX5 (XRD) which includes models of the tails and loops which are missing from the 4QLC structure. In this structure the tails are in an extended state away from the complex DNA, however in our experience they typically collapse onto the nucleosome DNA in an ensemble of states, which is in line with experimental results that show the tails are not freely exposed in solution. ^98^ For the purpose our study, the tails provide stability to the overall complex and do little to interact with linker histone. Lastly, experiments have shown the C-terminal tails of H2A and linker histones may have direct interactions with one another that facilitate binding. ^99–101^ However, because we are modeling only the globular portion of the linker histones that lack the C-terminal tails, we did not observe these interactions which may have an additional effect in shifting the binding equilibria between on- and off-dyad states.

Finally, Bednar *et al.* used cryo-EM and X-ray crystallography to show that the long C-terminal domain (∼ 100 residues) is oriented on the dyad and localized on a single linker DNA arm. In our off-dyad model (Figure 8), this tail would be aligned distal to the dyad and interacting along the outside of a single linker DNA. Here, the C-terminal linker histone tail could compete for binding with the long H3 tail along the same DNA arm potentially inducing a shift from the off- to on-dyad binding mode. The oppositely oriented N-terminal tail (∼ 20 to 100 residues) is much less studied and its role in off-dyad binding remains unclear. In our model, the N-terminal domain finds itself along the inside of the same DNA arm where it could interact with either linker DNA segment.

**Figure 8:**
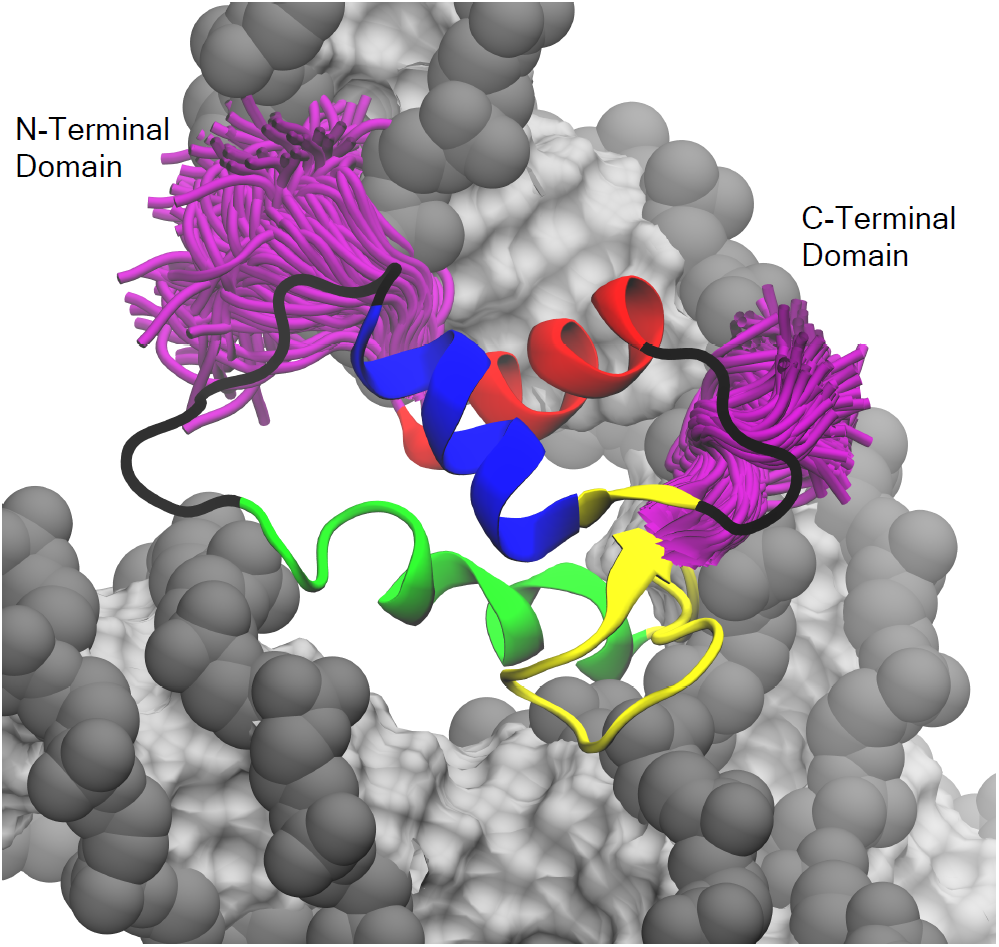
Example of terminal tail sampling in the off-dyad linker histones (genGH1 shown above). Shown in purple are the sampling of the terminal tails of every 100 frames from a single simulation trajectory. Colored by secondary structure: α-helix 1 (α1; red), α-helix 2 (α2; blue), α-helix 3 (α3; green), the β-sheet (yellow), and disordered regions (black). DNA is shown in gray (backbone) and silver (bases).

Overall, our study builds on notions of an ensemble of linker histone binding states within a highly dynamic chromatin fiber while emphasizing the contrasting influence of their variants on those structures and dynamics. Currently, the relative populations of these states within chromatin are still debatable. We subscribe to the hypothesis that the on- and off-dyad binding modes exist as an ensemble of states within chromatin fibers. ^47^ The relative populations of these states *in vivo* are likely due to a balance of not only the linker histone/nucleosome interactions examined here, but also factors outside the scope of this study such as DNA sequence, nucleosome repeat length, and the greater chromatin architecture. Potentially, coarse-grained models may be more adept at sampling the populations of binding states in mono- and poly-chromatosomes arrays. However, care should be taken in these models as the estimated binding energies calculated here demonstrate the importance of Van der Waals interactions within the chromatosome in addition to the more commonly considered electrostatic energies. Therefore, we emphasize that any model which attempts to recapitulate the physics underlying linker histone binding must carefully balance their electrostatic and Van der Waals components.

## Acknowledgments

The authors thank Dr. Francisco Rodríguez-Ropero for his assistance in preparing the off-dyad chromatosome system and Mr. Joseph Clayton for his assistance with many aspect of the simulation analysis. This work used the Extreme Science and Engineering Discovery Environment, which is supported by the National Science Foundation [ACI-1053575].

## References

[1] Kornberg, R. D. Chromatin structure: a repeating unit of histones and DNA. Science 1974, 184, 868–871.

[2] Olins, A. L.; Olins, D. E. Spheroid chromatin units (v bodies). Science 1974, 183, 330–332.

[3] Kaplan, N.; Moore, I. K.; Fondufe-Mittendorf, Y.; Gossett, A. J.; Tillo, D.; Field, Y.; LeProust, E. M.; Hughes, T. R.; Lieb, J. D.; Widom, J. et al. The DNA-encoded nucleosome organization of a eukaryotic genome. Nature 2009, 458, 362–366.

[4] Klemm, S. L.; Shipony, Z.; Greenleaf, W. J. Chromatin accessibility and the regulatory epigenome. Nat. Rev. Genet. 2019, 20, 207–220.

[5] Lai, W. K. M.; Pugh, B. F. Understanding nucleosome dynamics and their links to gene expression and DNA replication. Nat. Rev. Mol. Cell Biol. 2017, 18, 548–562.

[6] Koyama, M.; Kurumizaka, H. Structural diversity of the nucleosome. J. Biochem. 2018, 163, 85–95.

[7] Luger, K.; Mader, A. W.; Richmond, R. K.; Sargent, D. F.; Richmond, T. J. Crystal structure of the nucleosome core particle at 2.8 Å resolution. Nature 1997, 389, 251–260.

[8] McGinty, R. K.; Tan, S. Nucleosome structure and function. Chem. Rev. 2015, 115, 2255–2273.

[9] Cutter, A. R.; Hayes, J. J. A brief review of nucleosome structure. FEBS Lett. 2015, 589, 2914–2922.

[10] Zhou, K.; Gaullier, G.; Luger, K. Nucleosome structure and dynamics are coming of age. Nat. Struct. Mol. Biol. 2019, 26, 3–13.

[11] Noll, M.; Kornberg, R. D. Action of micrococcal nuclease on chromatin and the location of histone H1. J. Mol. Biol. 1977, 109, 393–404.

[12] Maresca, T. J.; Freedman, B. S.; Heald, R. Histone H1 is essential for mitotic chromosome architecture and segregation in Xenopus laevis egg extracts. J. Cell Biol. 2005, 169, 859–869.

[13] Routh, A.; Sandin, S.; Rhodes, D. Nucleosome repeat length and linker histone stoichiometry determine chromatin fiber structure. Proc. Natl. Acad. Sci. U.S.A. 2008, 105, 8872–8877.

[14] Hergeth, S. P.; Schneider, R. The H1 linker histones: multifunctional proteins beyond the nucleosomal core particle. EMBO Rep. 2015, 16, 1439–1453.

[15] Fyodorov, D. V.; Zhou, B. R.; Skoultchi, A. I.; Bai, Y. Emerging roles of linker histones in regulating chromatin structure and function. Nat. Rev. Mol. Cell Biol. 2018, 19, 192–206.

[16] Fan, Y.; Nikitina, T.; Zhao, J.; Fleury, T. J.; Bhattacharyya, R.; Bouhassira, E. E.; Stein, A.; Woodcock, C. L.; Skoultchi, A. I. Histone H1 depletion in mammals alters global chromatin structure but causes specific changes in gene regulation. Cell 2005, 123, 1199–1212.

[17] Shen, X.; Gorovsky, M. A. Linker histone H1 regulates specific gene expression but not global transcription in vivo. Cell 1996, 86, 475–483.

[18] Lu, X.; Wontakal, S. N.; Kavi, H.; Kim, B. J.; Guzzardo, P. M.; Emelyanov, A. V.; Xu, N.; Hannon, G. J.; Zavadil, J.; Fyodorov, D. V. et al. Drosophila H1 regulates the genetic activity of heterochromatin by recruitment of Su(var)3-9. Science 2013, 340, 78–81.

[19] Lee, H.; Habas, R.; Abate-Shen, C. MSX1 cooperates with histone H1b for inhibition of transcription and myogenesis. Science 2004, 304, 1675–1678.

[20] Zhang, Y.; Khan, D.; Delling, J.; Tobiasch, E. Mechanisms underlying the osteo- and adipo-differentiation of human mesenchymal stem cells. ScientificWorldJournal 2012, 2012, 793823.

[21] Christophorou, M. A.; Castelo-Branco, G.; Halley-Stott, R. P.; Oliveira, C. S.; Loos, R.; Radzisheuskaya, A.; Mowen, K. A.; Bertone, P.; Silva, J. C.; Zernicka-Goetz, M. et al. Citrullination regulates pluripotency and histone H1 binding to chromatin. Nature 2014, 507, 104–108.

[22] Thorslund, T.; Ripplinger, A.; Hoffmann, S.; Wild, T.; Uckelmann, M.; Villumsen, B.; Narita, T.; Sixma, T. K.; Choudhary, C.; Bekker-Jensen, S. et al. Histone H1 couples initiation and amplification of ubiquitin signalling after DNA damage. Nature 2015, 527, 389–393.

[23] Konishi, A.; Shimizu, S.; Hirota, J.; Takao, T.; Fan, Y.; Matsuoka, Y.; Zhang, L.; Yoneda, Y.; Fujii, Y.; Skoultchi, A. I. et al. Involvement of histone H1.2 in apoptosis induced by DNA double-strand breaks. Cell 2003, 114, 673–688.

[24] Lever, M. A.; Th’ng, J. P.; Sun, X.; Hendzel, M. J. Rapid exchange of histone H1.1 on chromatin in living human cells. Nature 2000, 408, 873–876.

[25] Misteli, T.; Gunjan, A.; Hock, R.; Bustin, M.; Brown, D. T. Dynamic binding of histone H1 to chromatin in living cells. Nature 2000, 408, 877–881.

[26] Catez, F.; Ueda, T.; Bustin, M. Determinants of histone H1 mobility and chromatin binding in living cells. Nat. Struct. Mol. Biol. 2006, 13, 305–310.

[27] Thoma, F.; Koller, T.; Klug, A. Involvement of histone H1 in the organization of the nucleosome and of the salt-dependent superstructures of chromatin. J. Cell Biol. 1979, 83, 403–427.

[28] Shimamura, A.; Sapp, M.; Rodriguez-Campos, A.; Worcel, A. Histone H1 represses transcription from minichromosomes assembled in vitro. Mol. Cell. Biol. 1989, 9, 5573–5584.

[29] Laybourn, P. J.; Kadonaga, J. T. Role of nucleosomal cores and histone H1 in regulation of transcription by RNA polymerase II. Science 1991, 254, 238–245.

[30] O’Neill, T. E.; Meersseman, G.; Pennings, S.; Bradbury, E. M. Deposition of histone H1 onto reconstituted nucleosome arrays inhibits both initiation and elongation of transcripts by T7 RNA polymerase. Nucleic Acids Res. 1995, 23, 1075–1082.

[31] Mishra, L. N.; Hayes, J. J. A nucleosome-free region locally abrogates histone H1-dependent restriction of linker DNA accessibility in chromatin. J. Biol. Chem. 2018, 293, 19191–19200.

[32] McGhee, J. D.; Felsenfeld, G. Nucleosome structure. Annu. Rev. Biochem. 1980, 49, 1115–1156.

[33] Zhou, B. R.; Jiang, J.; Feng, H.; Ghirlando, R.; Xiao, T. S.; Bai, Y. Structural Mechanisms of Nucleosome Recognition by Linker Histones. Mol. Cell 2015, 59, 628–638.

[34] Bednar, J.; Garcia-Saez, I.; Boopathi, R.; Cutter, A. R.; Papai, G.; Reymer, A.; Syed, S. H.; Lone, I. N.; Tonchev, O.; Crucifix, C. et al. Structure and Dynamics of a 197 bp Nucleosome in Complex with Linker Histone H1. Mol. Cell 2017, 66, 384–397.

[35] Syed, S. H.; Goutte-Gattat, D.; Becker, N.; Meyer, S.; Shukla, M. S.; Hayes, J. J.; Everaers, R.; Angelov, D.; Bednar, J.; Dimitrov, S. Single-base resolution mapping of H1-nucleosome interactions and 3D organization of the nucleosome. Proc. Natl. Acad. Sci. U.S.A. 2010, 107, 9620–9625.

[36] Zhou, B. R.; Feng, H.; Ghirlando, R.; Li, S.; Schwieters, C. D.; Bai, Y. A Small Number of Residues Can Determine if Linker Histones Are Bound On or Off Dyad in the Chromatosome. J. Mol. Biol. 2016, 428, 3948–3959.

[37] Song, F.; Chen, P.; Sun, D.; Wang, M.; Dong, L.; Liang, D.; Xu, R. M.; Zhu, P.; Li, G. Cryo-EM study of the chromatin fiber reveals a double helix twisted by tetranucleosomal units. Science 2014, 344, 376–380.

[38] Zhou, B. R.; Feng, H.; Kato, H.; Dai, L.; Yang, Y.; Zhou, Y.; Bai, Y. Structural insights into the histone H1-nucleosome complex. Proc. Natl. Acad. Sci. U.S.A. 2013, 110, 19390–19395.

[39] Stehr, R.; Kepper, N.; Rippe, K.; Wedemann, G. The effect of internucleosomal interaction on folding of the chromatin fiber. Biophys. J. 2008, 95, 3677–3691.

[40] Kepper, N.; Foethke, D.; Stehr, R.; Wedemann, G.; Rippe, K. Nucleosome geometry and internucleosomal interactions control the chromatin fiber conformation. Biophys. J. 2008, 95, 3692–3705.

[41] Perišić, O.; Portillo-Ledesma, S.; Schlick, T. Sensitive effect of linker histone binding mode and subtype on chromatin condensation. Nucleic Acids Res. 2019,

[42] Garcia-Saez, I.; Menoni, H.; Boopathi, R.; Shukla, M. S.; Soueidan, L.; Noirclerc-Savoye, M.; Le Roy, A.; Skoufias, D. A.; Bednar, J.; Hamiche, A. et al. Structure of an H1-Bound 6-Nucleosome Array Reveals an Untwisted Two-Start Chromatin Fiber Conformation. Mol. Cell 2018, 72, 902–915.

[43] Szerlong, H. J.; Hansen, J. C. Nucleosome distribution and linker DNA: connecting nuclear function to dynamic chromatin structure. Biochem. Cell Biol. 2011, 89, 24–34.

[44] Grigoryev, S. A. Nucleosome spacing and chromatin higher-order folding. Nucleus 2012, 3, 493–499.

[45] Grigoryev, S. A.; Schubert, M. Unraveling the multiplex folding of nucleosome chains in higher order chromatin. Essays Biochem. 2019, 63, 109–121.

[46] Zhou, B. R.; Jiang, J.; Ghirlando, R.; Norouzi, D.; Sathish Yadav, K. N.; Feng, H.; Wang, R.; Zhang, P.; Zhurkin, V.; Bai, Y. Revisit of Reconstituted 30-nm Nucleosome Arrays Reveals an Ensemble of Dynamic Structures. J. Mol. Biol. 2018, 430, 3093–3110.

[47] Öztürk, M. A.; Cojocaru, V.; Wade, R. C. Toward an Ensemble View of Chromatosome Structure: A Paradigm Shift from One to Many. Structure 2018, 26, 1050–1057.

[48] Talbert, P. B.; Ahmad, K.; Almouzni, G.; Ausio, J.; Berger, F.; Bhalla, P. L.; Bonner, W. M.; Cande, W. Z.; Chadwick, B. P.; Chan, S. W. et al. A unified phylogeny-based nomenclature for histone variants. Epigenetics Chromatin 2012, 5, 7.

[49] Draizen, E. J.; Shaytan, A. K.; Marino-Ramirez, L.; Talbert, P. B.; Landsman, D.; Panchenko, A. R. HistoneDB 2.0: a histone database with variants–an integrated resource to explore histones and their variants. Database (Oxford) 2016, 2016.

[50] Bascom, G. D.; Schlick, T. Chromatin Fiber Folding Directed by Cooperative Histone Tail Acetylation and Linker Histone Binding. Biophys. J. 2018, 114, 2376–2385.

[51] Luque, A.; Ozer, G.; Schlick, T. Correlation among DNA Linker Length, Linker Histone Concentration, and Histone Tails in Chromatin. Biophys. J. 2016, 110, 2309–2319.

[52] Nizovtseva, E. V.; Clauvelin, N.; Todolli, S.; Polikanov, Y. S.; Kulaeva, O. I.; Wengrzynek, S.; Olson, W. K.; Studitsky, V. M. Nucleosome-free DNA regions differentially affect distant communication in chromatin. Nucleic Acids Res. 2017, 45, 3059–3067.

[53] Öztürk, M. A.; Pachov, G. V.; Wade, R. C.; Cojocaru, V. Conformational selection and dynamic adaptation upon linker histone binding to the nucleosome. Nucleic Acids Res. 2016, 44, 6599–6613.

[54] Öztürk, M. A.; Cojocaru, V.; Wade, R. C. Dependence of Chromatosome Structure on Linker Histone Sequence and Posttranslational Modification. Biophys. J. 2018, 114, 2363–2375.

[55] Ramakrishnan, V.; Finch, J. T.; Graziano, V.; Lee, P. L.; Sweet, R. M. Crystal structure of globular domain of histone H5 and its implications for nucleosome binding. Nature 1993, 362, 219–223.

[56] Pachov, G. V.; Gabdoulline, R. R.; Wade, R. C. On the structure and dynamics of the complex of the nucleosome and the linker histone. Nucleic Acids Res. 2011, 39, 5255–5263.

[57] Brown, D. T.; Izard, T.; Misteli, T. Mapping the interaction surface of linker histone H1(0) with the nucleosome of native chromatin in vivo. Nat. Struct. Mol. Biol. 2006, 13, 250–255.

[58] Bharath, M. M.; Chandra, N. R.; Rao, M. R. Molecular modeling of the chromatosome particle. Nucleic Acids Res. 2003, 31, 4264–4274.

[59] Zhou, Y. B.; Gerchman, S. E.; Ramakrishnan, V.; Travers, A.; Muyldermans, S. Position and orientation of the globular domain of linker histone H5 on the nucleosome. Nature 1998, 395, 402–405.

[60] George, E. M.; Izard, T.; Anderson, S. D.; Brown, D. T. Nucleosome interaction surface of linker histone H1c is distinct from that of H1(0). J. Biol. Chem. 2010, 285, 20891–20896.

[61] Cutter, A. R.; Hayes, J. J. Linker histones: novel insights into structure-specific recognition of the nucleosome. Biochem. Cell Biol. 2017, 95, 171–178.

[62] Davey, C. A.; Sargent, D. F.; Luger, K.; Maeder, A. W.; Richmond, T. J. Solvent mediated interactions in the structure of the nucleosome core particle at 1.9 a resolution. J. Mol. Biol. 2002, 319, 1097–1113.

[63] Dorigo, B.; Schalch, T.; Kulangara, A.; Duda, S.; Schroeder, R. R.; Richmond, T. J. Nucleosome arrays reveal the two-start organization of the chromatin fiber. Science 2004, 306, 1571–1573.

[64] Thastrom, A.; Bingham, L. M.; Widom, J. Nucleosomal locations of dominant DNA sequence motifs for histone-DNA interactions and nucleosome positioning. J. Mol. Biol. 2004, 338, 695–709.

[65] Sali, A.; Blundell, T. L. Comparative protein modelling by satisfaction of spatial restraints. J. Mol. Biol. 1993, 234, 779–815.

[66] Pettersen, E. F.; Goddard, T. D.; Huang, C. C.; Couch, G. S.; Greenblatt, D. M.; Meng, E. C.; Ferrin, T. E. UCSF Chimera–a visualization system for exploratory research and analysis. J Comput Chem 2004, 25, 1605–1612.

[67] Wriggers, W. Conventions and workflows for using Situs. Acta Crystallogr. D Biol. Crystallogr. 2012, 68, 344–351.

[68] Wriggers, W.; Milligan, R. A.; McCammon, J. A. Situs: A package for docking crystal structures into low-resolution maps from electron microscopy. J. Struct. Biol. 1999, 125, 185–195.

[69] Lopez-Blanco, J. R.; Garzon, J. I.; Chacon, P. iMod: multipurpose normal mode analysis in internal coordinates. Bioinformatics 2011, 27, 2843–2850.

[70] Case, D. A.; Betz, R. M.; Cerutti, D. S.; Cheatham, T. E.; Darden, T. A.; Duke, R. E.; Giese, T. J.; Gohlke, H.; Goetz, A. W.; Homeyer, N. et al. AMBER 2016; University of California, San Francisco, 2016.

[71] Jorgensen, W. L.; Chandrasekhar, J.; Madura, J. D.; Impey, R. W.; Klein, M. L. Comparison of simple potential functions for simulating liquid water. J Chem Phys 1983, 79, 926–935.

[72] Mahoney, M. W.; Jorgensen, W. L. A five-site model for liquid water and the reproduction of the density anomaly by rigid, nonpolarizable potential functions. J Chem Phys 1983, 112, 8910–8922.

[73] Joung, I. S.; Cheatham, T. E. Determination of alkali and halide monovalent ion parameters for use in explicitly solvated biomolecular simulations. J Phys Chem B 2008, 112, 9020–9041.

[74] Joung, I. S.; Cheatham, T. E. Molecular dynamics simulations of the dynamic and energetic properties of alkali and halide ions using water-model-specific ion parameters. J Phys Chem B 2009, 113, 13279–13290.

[75] Maier, J. A.; Martinez, C.; Kasavajhala, K.; Wickstrom, L.; Hauser, K. E.; Simmerling, C. ff14SB: Improving the Accuracy of Protein Side Chain and Backbone Parameters from ff99SB. J Chem Theory Comput 2015, 11, 3696–3713.

[76] Ivani, I.; Dans, P. D.; Noy, A.; Perez, A.; Faustino, I.; Hospital, A.; Walther, J.; Andrio, P.; Goni, R.; Balaceanu, A. et al. Parmbsc1: a refined force field for DNA simulations. Nat. Methods 2016, 13, 55–58.

[77] Phillips, J. C.; Braun, R.; Wang, W.; Gumbart, J.; Tajkhorshid, E.; Villa, E.; Chipot, C.; Skeel, R. D.; Kale, L.; Schulten, K. Scalable molecular dynamics with NAMD. J Comput Chem 2005, 26, 1781–1802.

[78] Darden, T.; York, D.; Pedersen, L. Particle mesh Ewald: An Nlog(N) method for Ewald sums in large systems. J. Chem. Phys. 1993, 98, 10089–10092.

[79] Hoover, W. G. Canonical dynamics: Equilibrium phase-space distributions. Phys Rev A Gen Phys 1985, 31, 1695–1697.

[80] Towns, J.; Cockerill, T.; Dahan, M.; Foster, I.; Gaither, K.; Grimshaw, A.; Hazlewood, V.; Lathrop, S.; Lifka, D.; Peterson, G. et al. XSEDE: accelerating scientific discovery. Computing in Science & Engineering 2014, 16, 62–74.

[81] Henin, J.; Fiorin, G.; Chipot, C.; Klein, M. L. Exploring Multidimensional Free Energy Landscapes Using Time-Dependent Biases on Collective Variables. J Chem Theory Comput 2010, 6, 35–47.

[82] Kumar, S.; Rosenberg, J. M.; Bouzida, D.; Swendsen, R. H.; Kollman, P. A. THE weighted histogram analysis method for free-energy calculations on biomolecules. I. The method. Journal of Computational Chemistry 1992, 13, 1011–1021.

[83] Grossfield, A. WHAM: the weighted histogram analysis method.

[84] Humphrey, W.; Dalke, A.; Schulten, K. VMD: visual molecular dynamics. J Mol Graph 1996, 14, 33–38.

[85] Miller, B. R.; McGee, T. D.; Swails, J. M.; Homeyer, N.; Gohlke, H.; Roitberg, A. E. MMPBSA.py: An Efficient Program for End-State Free Energy Calculations. J Chem Theory Comput 2012, 8, 3314–3321.

[86] Nguyen, H.; Perez, A.; Bermeo, S.; Simmerling, C. Refinement of Generalized Born Implicit Solvation Parameters for Nucleic Acids and Their Complexes with Proteins. J Chem Theory Comput 2015, 11, 3714–3728.

[87] Johnson, S. C. Hierarchical clustering schemes. Psychometrika 1967, 32, 241–254.

[88] Kvålseth, T. O. On Normalized Mutual Information: Measure Derivations and Properties. Entropy 2017, 631, 1–14.

[89] McClendon, C. L.; Hua, L.; Barreiro, A.; Jacobson, M. P. Comparing Conformational Ensembles Using the Kullback-Leibler Divergence Expansion. J Chem Theory Comput 2012, 8, 2115–2126.

[90] Piana, S.; Donchev, A. G.; Robustelli, P.; Shaw, D. E. Water dispersion interactions strongly influence simulated structural properties of disordered protein states. J Phys Chem B 2015, 119, 5113–5123.

[91] Battiste, J. L.; Wagner, G. Utilization of site-directed spin labeling and high-resolution heteronuclear nuclear magnetic resonance for global fold determination of large proteins with limited nuclear overhauser effect data. Biochemistry 2000, 39, 5355–5365.

[92] Clore, G. M. Practical Aspects of Paramagnetic Relaxation Enhancement in Biological Macromolecules. Meth. Enzymol. 2015, 564, 485–497.

[93] Shaytan, A. K.; Armeev, G. A.; Goncearenco, A.; Zhurkin, V. B.; Landsman, D.; Panchenko, A. R. Coupling between Histone Conformations and DNA Geometry in Nucleosomes on a Microsecond Timescale: Atomistic Insights into Nucleosome Functions. J. Mol. Biol. 2016, 428, 221–237.

[94] Miller, B. R.; McGee, T. D.; Swails, J. M.; Homeyer, N.; Gohlke, H.; Roitberg, A. E. MMPBSA.py: An Efficient Program for End-State Free Energy Calculations. J Chem Theory Comput 2012, 8, 3314–3321.

[95] Hou, T.; Wang, J.; Li, Y.; Wang, W. Assessing the performance of the MM/PBSA and MM/GBSA methods. 1. The accuracy of binding free energy calculations based on molecular dynamics simulations. J Chem Inf Model 2011, 51, 69–82.

[96] Genheden, S.; Ryde, U. The MM/PBSA and MM/GBSA methods to estimate ligand-binding affinities. Expert Opin Drug Discov 2015, 10, 449–461.

[97] Genheden, S.; Ryde, U. Will molecular dynamics simulations of proteins ever reach equilibrium? Phys Chem Chem Phys 2012, 14, 8662–8677.

[98] Morrison, E. A.; Bowerman, S.; Sylvers, K. L.; Wereszczynski, J.; Musselman, C. A. The conformation of the histone H3 tail inhibits association of the BPTF PHD finger with the nucleosome. Elife 2018, 7.

[99] Jason, L. J. M.; Finn, R. M.; Lindsey, G.; Ausió, J. Histone H2A ubiquitination does not preclude histone H1 binding, but it facilitates its association with the nucleosome. J Biol Chem 2005, 280, 4975–4982.

[100] Vogler, C.; Huber, C.; Waldmann, T.; Ettig, R.; Braun, L.; Izzo, A.; Daujat, S.; Chassignet, I.; Lopez-Contreras, A. J.; Fernandez-Capetillo, O. et al. Histone H2A C-terminus regulates chromatin dynamics, remodeling, and histone H1 binding. PLoS Genet. 2010, 6, e1001234.

[101] Shukla, M. S.; Syed, S. H.; Goutte-Gattat, D.; Richard, J. L.; Montel, F.; Hamiche, A.; Travers, A.; Faivre-Moskalenko, C.; Bednar, J.; Hayes, J. J. et al. The docking domain of histone H2A is required for H1 binding and RSC-mediated nucleosome remodeling. Nucleic Acids Res. 2011, 39, 2559–2570.

