## Supplemental Information for "Elucidating the Influence of Linker Histone Variants on Chromatosome Dynamics and Energetics"

### Supporting Information for Elucidating the Influence of Linker Histone variants on Chromatosome Dynamics and Energetic

### S1 Normalized Mutual Information

The mutual information (MI) between angles was calculated to determine the correlations between each pair of angles by:[88]

$$I\left(x,y\right)=\sum_{i=1}^{I}\sum_{j=1}^{J}p\left(x_i,y_j\right)\log\left(\frac{p\left(x_i,y_j\right)}{p\left(x_i\right)p\left(y_j\right)}\right) \tag{1}$$

where $I$ and $J$ represent the number of bins in the histograms of the two angles to be compared, $x$ and $y$. The normalized mutual information (NMI), $I^*$, was computed by:

$$H_x=\sum_{i=1}^{I}-p\left(x_i\right)\log p\left(x_i\right) \tag{2a}$$

$$H_y=\sum_{j=1}^{J}-p\left(y_j\right)\log p\left(y_j\right) \tag{2b}$$

$$I^*=\frac{I\left(x,y\right)}{\min\left(H_x,H_y\right)} \tag{2c}$$

where $I^*$ has the advantageous property that all values are in the range of 0 to 1 and are therefore easier to interpret than standard mutual information (MI).

### S2 Description of Linker DNA Angle Calculation

Below is a description of how we calculated the alpha (in-plane motions) and beta (out-of-plane motions) angles used to describe the linker DNA dynamics in the main text of the manuscript.

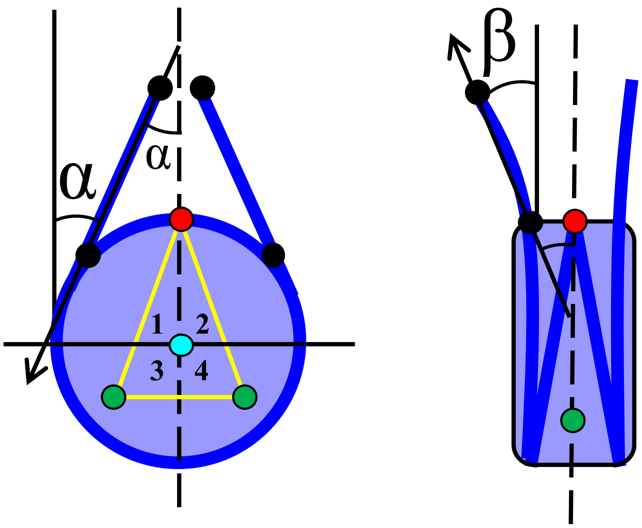


Figure S1: Diagram of the points used to construct vectors which were used to calculate the alpha (left) and beta (right) angles in Figures 6, S15, and S16

#### S2.1 The Black Points

These points create the vectors used to calculate both the alpha and beta angles. They are the midpoints of the C1' atoms between the base pairs at the origin (BP 10-325 and BP 158-177) and terminus (BP 1-334 and BP 167-168) of the exit and entry linker DNA. We call the line connecting these points, the Linker Vector.

#### S2.2 Alpha Angles

The Linker Vector intersects with the Dyad Vector to create the alpha angle (α). The Dyad Vector is the line connected by the dyad point (C1' midpoint of BP 83-250) and the histone core center-of-mass, which is defined as the alpha-carbon center of mass of all alpha-helical residues. We only consider residues within an alpha helix to reduce noise.

#### S2.3 Beta Angles

The beta angle is defined as the angle of the Linker Vector intersecting with nucleosomal plane. The nucleosomal plane is defined by three points. The first is the dyad point, defined above and represented as the red point in Figure S1. The next two points are green in Figure S1. For reference, we split the nucleosome into four quadrants. The green points are the C1' center-of-mass of the DNA in quadrants three and four because we believe the portion of DNA proximal to the linker DNA to be the most rigid.

### S3 Supplemental Tables

Table S1: Binding free-energy estimates from MM/GBSA calculations for GH1.0 and genGH1 in on and off-dyad states. $\Delta E_{total}$ represents the total MM/GBSA binding energy estimate. $\Delta E_{internal}$ is the internal energy change and is the sum of the change in bond, angle, and dihedral energy changes ($\Delta E_{bonds}$, $\Delta E_{angles}$, and $\Delta E_{dihedrals}$), $\Delta E_{elec}$ is the sum of the molecular mechanics electrostatics as well as the Generalized Born electrostatic terms ( $\Delta E_{elect\_MM}$ and $\Delta E_{solv\_GB}$) and $\Delta E_{VdW}$ is the sum of the molecular mechanics Van der Waals and Generalized Born nonpolar terms ($\Delta E_{solv\_GB}$ and $\Delta E_{solv\_SA}$). All energies are in kcal/mol.

| Linker Histone | Position | $\Delta E_{total}$ | $\Delta E_{internal}$ | $\Delta E_{elec}$ | $\Delta E_{VdW}$ |
|---|---|---|---|---|---|
| genGH1 | On-dyad | -174.6 ± 37.0 | -18.1 ±32.5 | -59.5 ±27.2 | -97.0 ±23.0 |
| | Off-dyad | -84.7 ±37.8 | 10.9 ±33.0 | 33.4 ±28.3 | -129.0 ±23.6 |
| GH1.0 | On-dyad | -221.6 ± 37.8 | -13.4 ±32.2 | -61.9 ±24.7 | -146.4 ±24.4 |
| | Off-dyad | -58.3 ± 36.5 | -10.1 ± 33.1 | 7.1 ± 24.2 | -55.4 ± 22.0 |

| Linker Histone | Position | $\Delta E_{total}$ | $\Delta E_{bonds}$ | $\Delta E_{angles}$ | $\Delta E_{dihedrals}$ | $\Delta E_{elect\_MM}$ | $\Delta E_{VdW\_MM}$ | $\Delta E_{solv\_GB}$ | $\Delta E_{solv\_SA}$ |
|---|---|---|---|---|---|---|---|---|---|
| genGH1 | On-dyad | -174.6 ± 37.0 | -1.3 ± 17.3 | -10.3 ± 24.8 | -6.6 ± 15.1 | -16575.4 ± 315.1 | -86.4 ± 22.3 | 16515.8 ± 318.5 | -10.6 ± 2.3 |
| | Off-dyad | -84.7 ±37.8 | 0.8 ± 17.4 | 18.7 ± 24.9 | -8.6 ± 15.2 | -15219.8 ± 422.7 | -114.2 ± 22.3 | 15253.2 ± 420.1 | -14.8 ± 2.1 |
| GH1.0 | On-dyad | -221.6 ± 37.8 | -3.5 ± 17.2 | -12.4 ±24.5 | 2.5 ± 15.0 | -15635.6 ±302.9 | -129.8 ±22.9 | 15573.8 ± 303.1 | -16.5 ± 2.2 |
| | Off-dyad | -58.3 ± 36.5 | 0.8 ± 17.0 | -2.9 ± 25.0 | -8.0 ± 15.1 | -14771.6 ± 328.7 | -50.4 ± 20.8 | 14778.7 ± 327.2 | -4.9 ± 1.9 |

Table S2: Kullback-Leibler divergence values for one dimensional probability distributions of DNA

| Linker Histone | Position | α-Entry | β-Entry | α-Exit | β-Exit |
|---|---|---|---|---|---|
| genGH1 | On-Dyad | 2.23 | 1.82 | 1.22 | 3.79 |
| | Off-Dyad | 0.37 | 0.31 | 0.22 | 6.34 |
| GH1.0 | On-Dyad | 2.61 | 1.25 | 2.42 | 1.03 |
| | Off-Dyad | 0.08 | 0.55 | 3.26 | 1.46 |

Table S3: Corresponding residue numbers for the points in Figure S2. Only the backbone atoms were considered for each residue.

| | genGH1 | | GH1.0 | |
|---|---|---|---|---|
| Point | Canonical Number | Res. | Canonical Number | Res. |
| A | 89 | PHE | 68 | ILE |
| | 90 | ILE | 69 | LYS |
| | 91 | LYS | 70 | LEU |
| | 92 | LYS | 71 | SER |
| B | 93 | TYR | 72 | ILE |
| | 94 | LEU | 73 | ARG |
| | 95 | LYS | 74 | ARG |
| | 96 | SER | 75 | LEU |
| C | 107 | LYS | 85 | LYS |
| | 108 | GLY | 86 | GLY |
| | 109 | LYS | 87 | VAL |
| | 110 | GLY | 88 | GLY |
| D | 102 | LYS | 80 | VAL |
| | 103 | LEU | 81 | LEU |
| | 113 | GLY | 91 | GLY |
| | 114 | SER | 92 | SER |

#### S4 Supplemental Figures

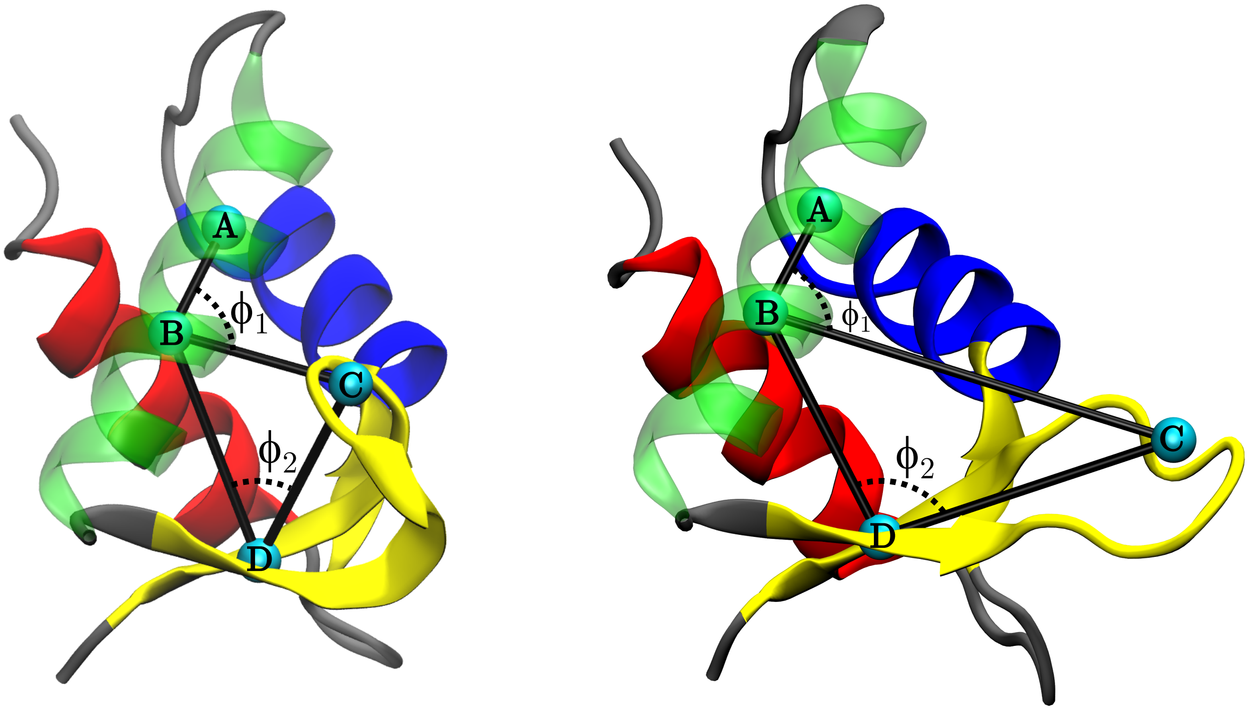


Figure S2: Diagrams for the angles, $\phi_1$ and $\phi_2$ (deg), used in the linker histone analyses. The points, seen above as light blue spheres, were defined briefly as follows using the backbone atoms of each residue: A) The geometric center of the second helical turn of the $\alpha$3 helix (green), B) the geometric center of the third helical turn of the $\alpha$3 helix, C) the geometric center of the $\beta$-loop from the closed-state, and D) the geometric center of the two residues in the center of each $\beta$-strand (four total residues). For more detail see Table S3.

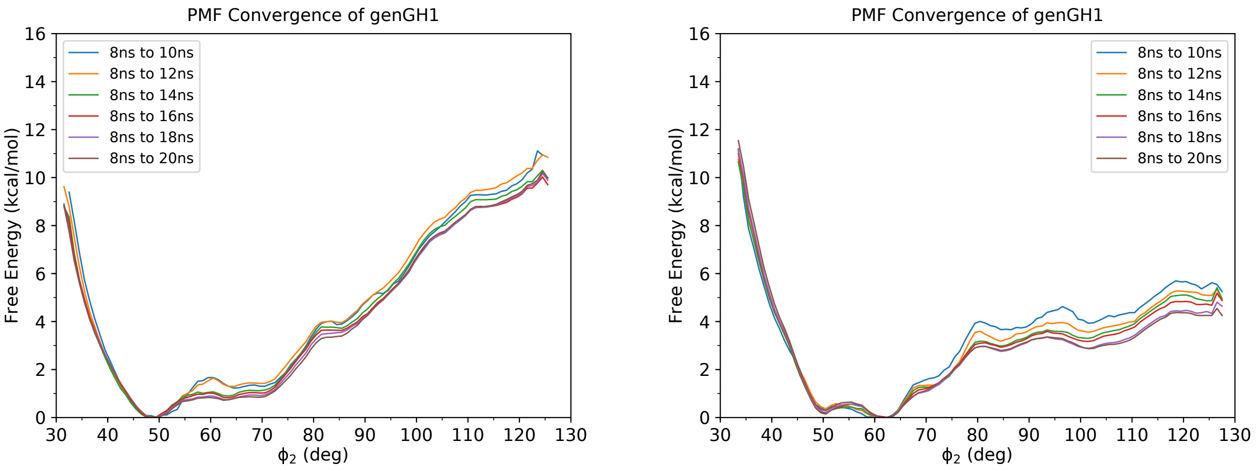


Figure S3: To depict convergence, shown is the change in PMFs from the umbrella sampling simulations through time.

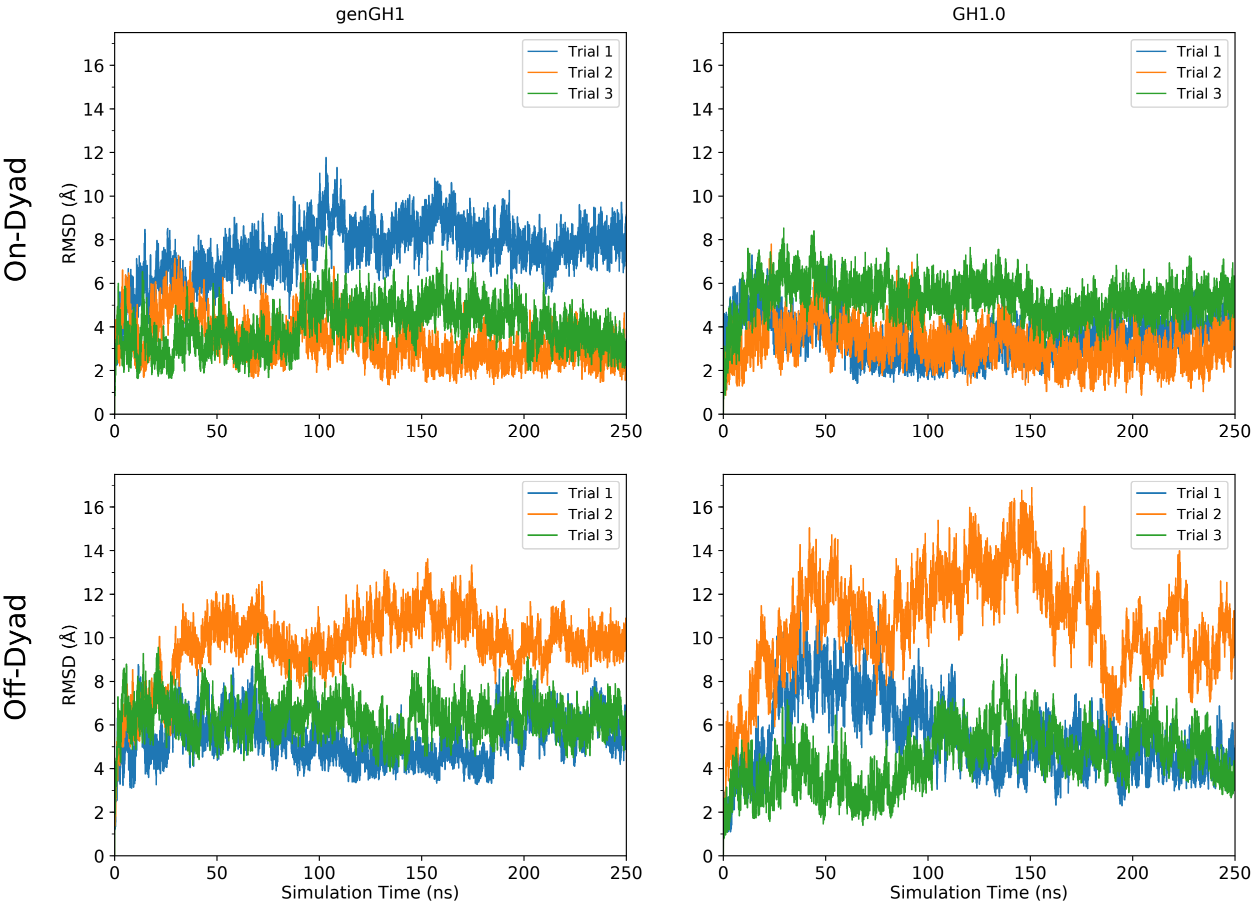


Figure S4: Root-mean-square deviation (RMSD) of the linker histone backbone atoms with respect to the alpha helical backbone atoms of the core histones. Shown are linker histones genGH1 and GH1.0 in both the on- and off-dyad binding modes. Each line represents a separate simulation totalling in 750 ns of simulation per system.

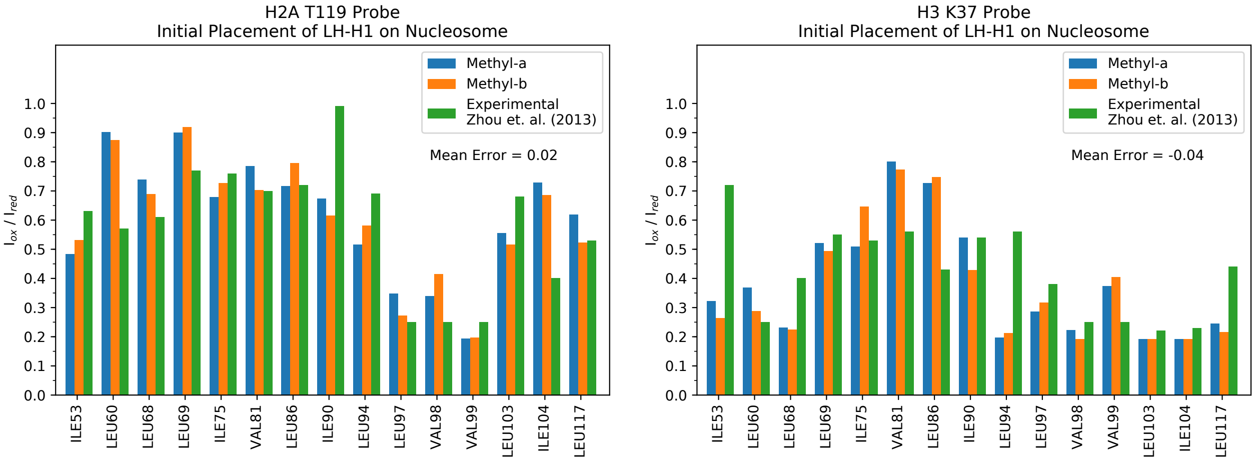


Figure S5: Theoretical paramagnetic relaxation enhancement intensity ratios ($I_{ox}/I_{red}$) for the initial placement of the linker histone genGH1 onto the extended nucleosome core particle for probe residues H2A T119 and H3 K37. Each residue contains two terminal methyl groups labeled methyl-a (blue) and methyl-b (orange), while experimental results (green) show only a solitary signal. The mean error is given in the top right-hand corner of each graph.

#### Comparison of Off-Dyad Models

Comparison of the off-dyad model used in this study with the model from Zhou *et al.* Figure S6 shows that the two are similar and differ only by a rotation of the linker histone. Comparison of the number of the linker histone/DNA contacts between the systems show that our system has increased stabilizing interactions, suggesting an energetically more favorable initial conformation (Figures S7-S9).

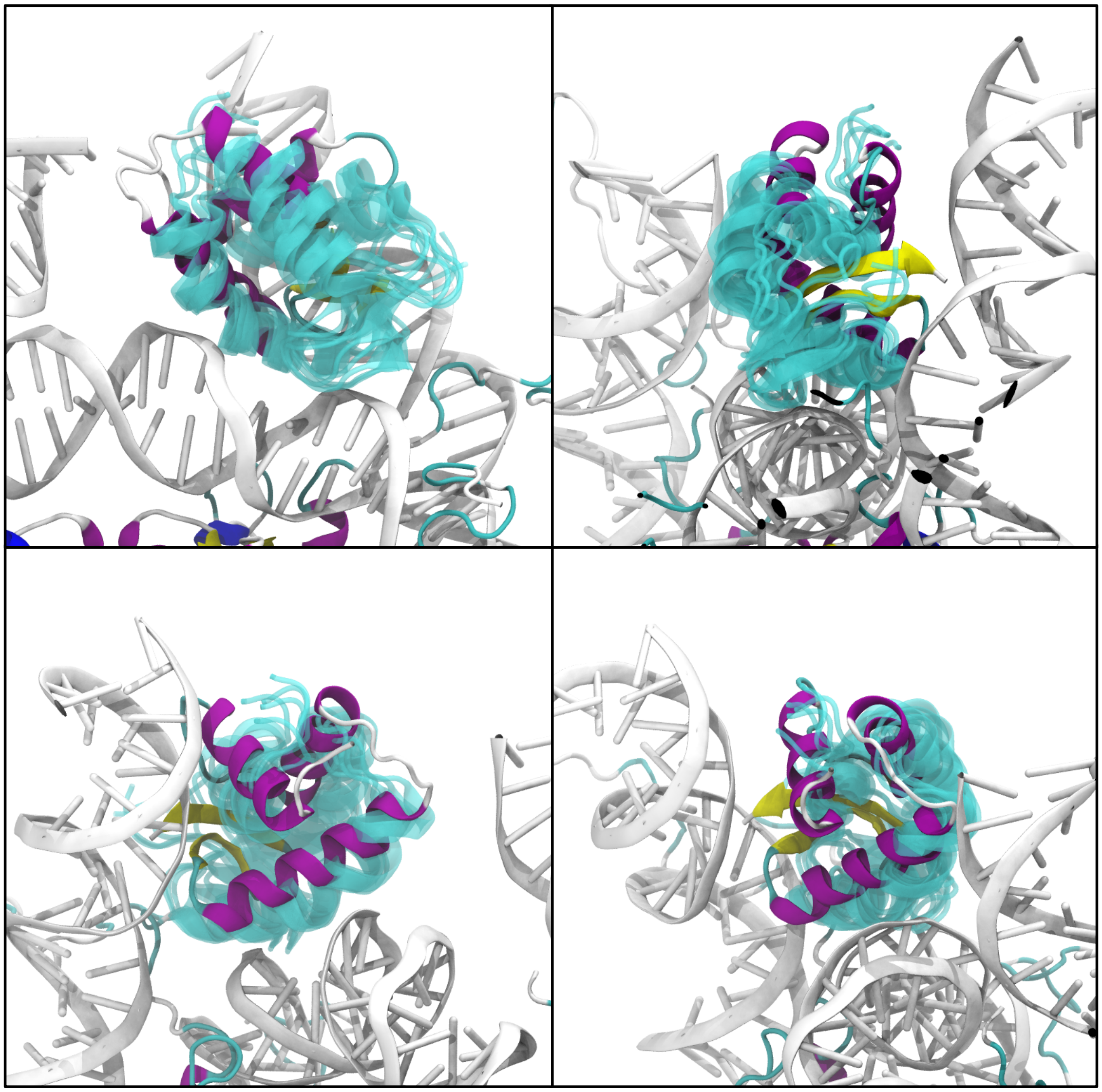

Figure S6: Shown is a physical comparison of binding pose in the Zhou *et al.*[38] off-dyad model (purple) and the clustered binding states sampled by the genGH1 used in throughout this study (cyan). The DNA can be seen in white.

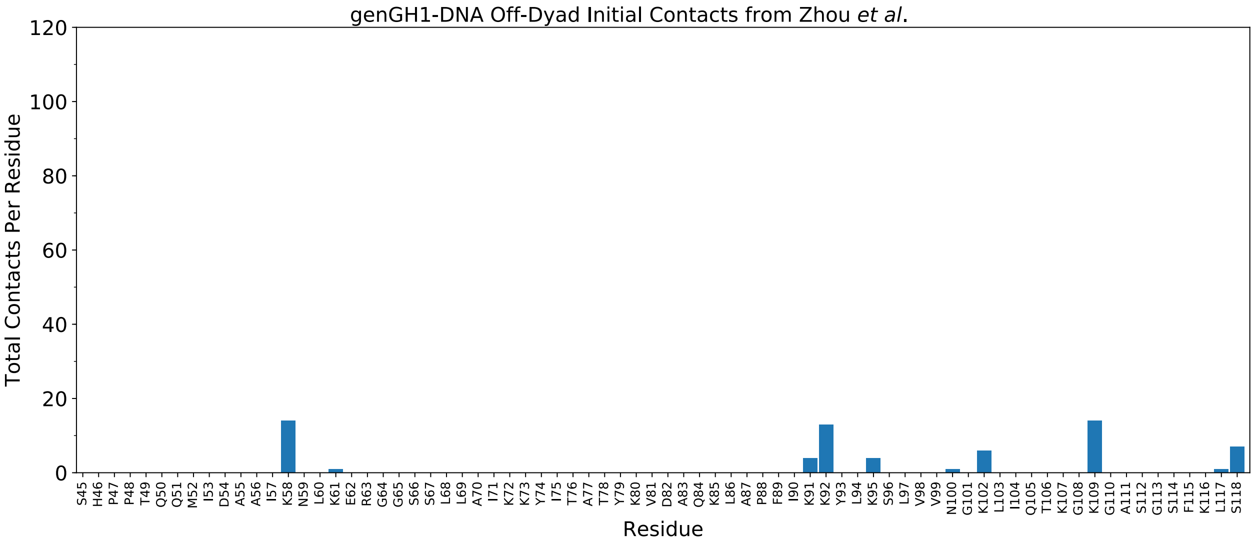


Figure S7: Initial residue contacts with DNA and the off-dyad genGH1 model from Zhou *et al.*[38]

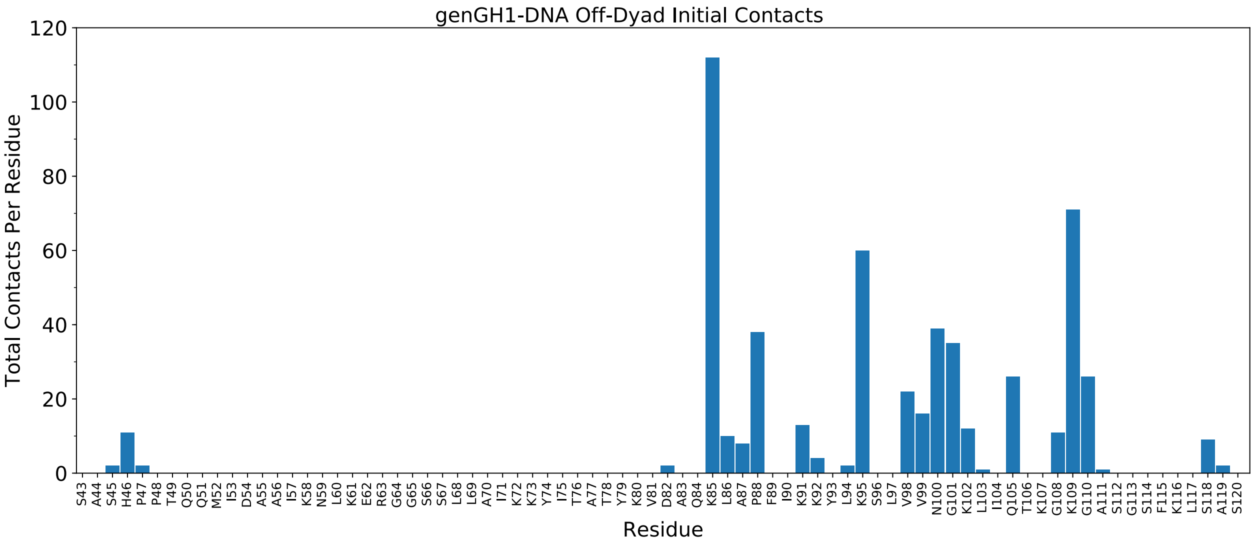


Figure S8: Initial residue contacts with DNA and the off-dyad genGH1 prior to running simulations.

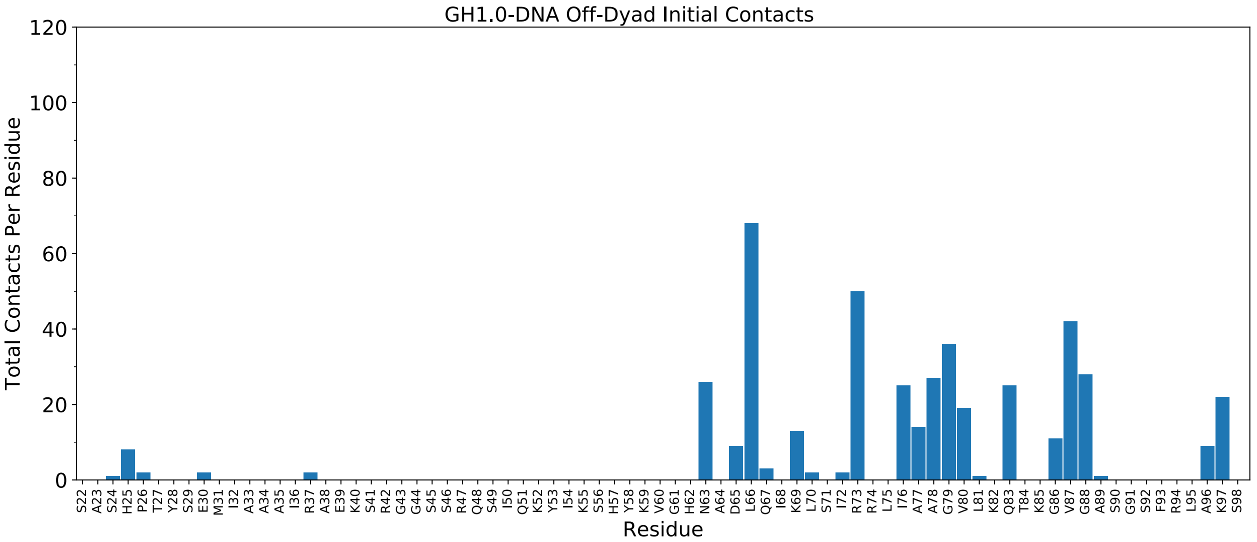


Figure S9: Initial residue contacts with DNA and the off-dyad GH1.0 prior to running simulations.

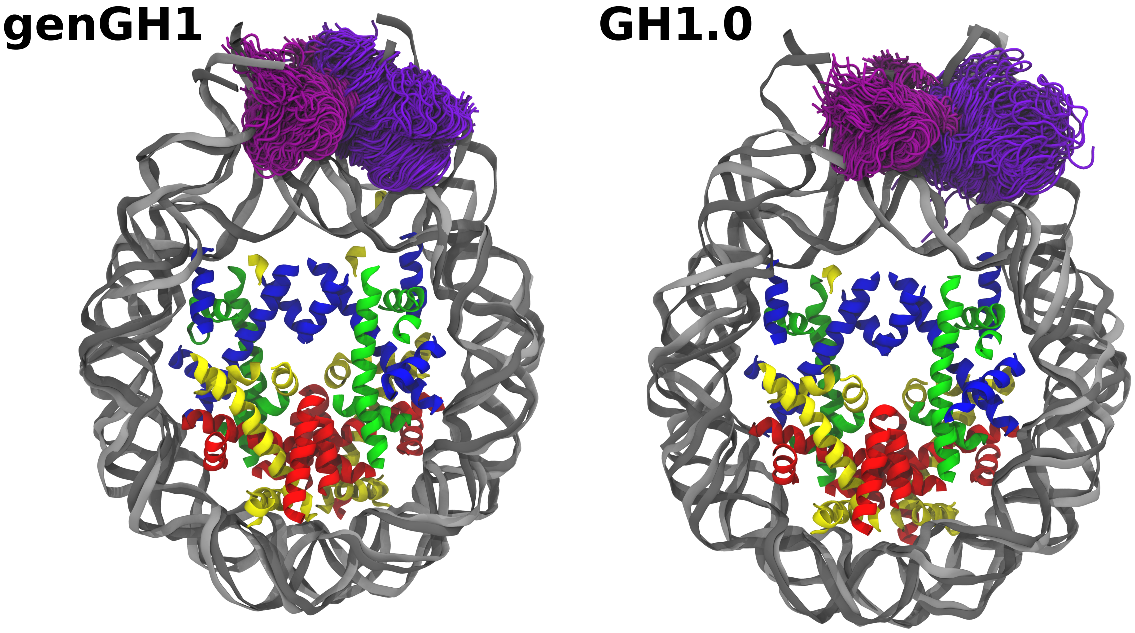


Figure S10: Shown are the distributions of genGH1 (left) and GH1.0 (right) in both the on- and off-dyad binding modes (lighter purple and darker violet, respectively). The DNA backbone is shown as gray (on-dyad) and silver (off-dyad). Core histones are shown without their tails and loops for the sake of clarity.

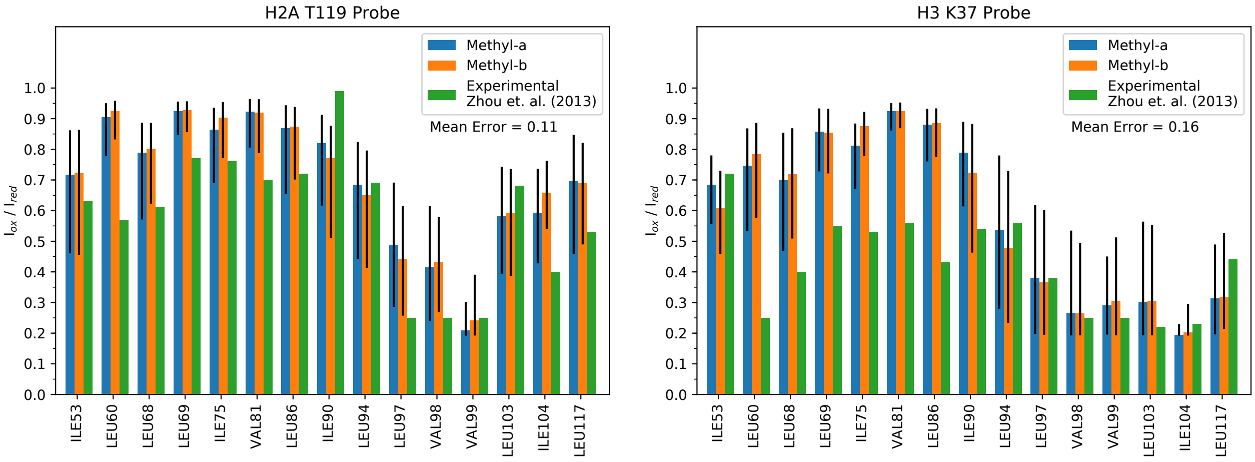


Figure S11: Time averaged theoretical paramagnetic relaxation enhancement intensity ratios ($I_{ox}/I_{red}$) over the equilibrated portions of the MD trajectories of the linker histone genGH1 onto the extended nucleosome core particle for probe residues H2A T119 and H3 K37. Each residue contains two terminal methyl groups labeled methyl-a (blue) and methyl-b (orange), while experimental results (green) show only a solitary signal. The mean error is given in the top right-hand corner of each graph.

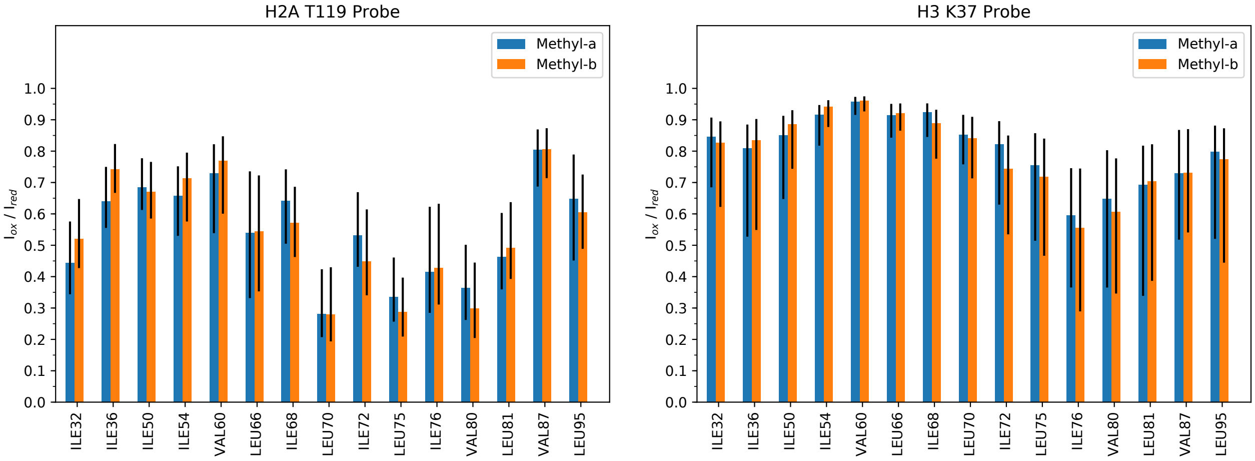


Figure S12: Theoretical paramagnetic relaxation enhancement ratios for the time-averaged MD simulations of GH1.0 on the extended nucleosome core particle.

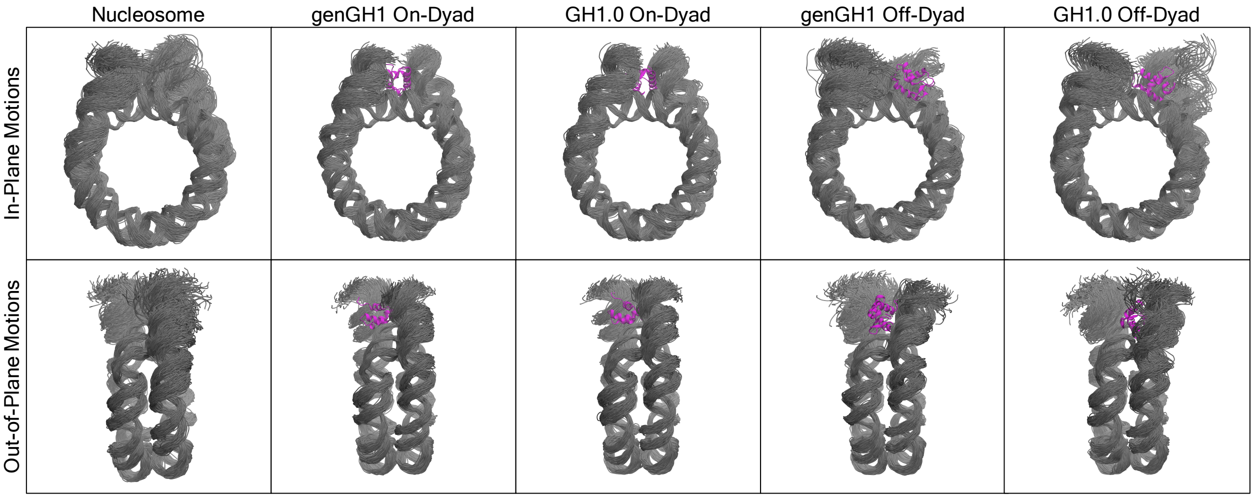


Figure S13: In-plane (top) and out-of-plane (bottom) sampling of the DNA throughout the length of the equilibrated systems. Shown are 150 representative frames of DNA (grey) along with the linker histone (purple) from a single frame for reference. Figures inspired by work from Shaytan *et al.*[93]

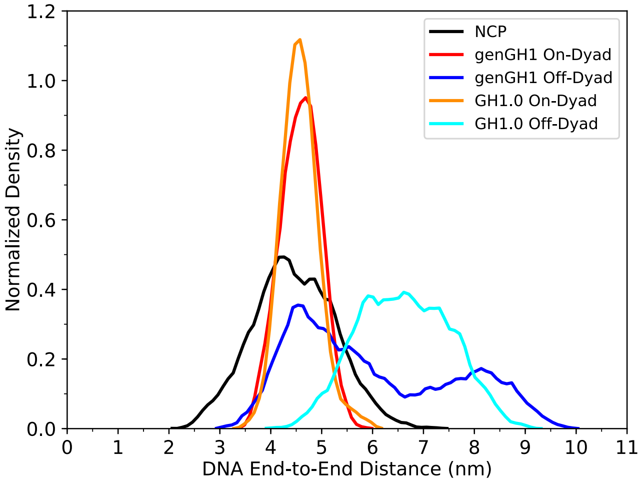


Figure S14: Normalized histograms for the end-to-end distances between the center-of-mass of the terminal base pairs of each linker DNA arm.

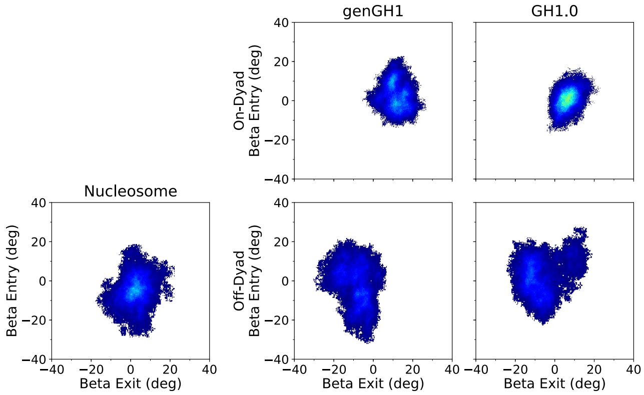


Figure S15: Density plots of beta linker DNA angles. Each plot shows a 2D histogram of the entry versus the exit beta linker DNA angles. Density ranges from dark blue (lowest) to red (highest). Each plot contains the linker DNA from the nucleosome core particle along with genGH1 and GH1.0 in the on- and off-dyad binding modes.

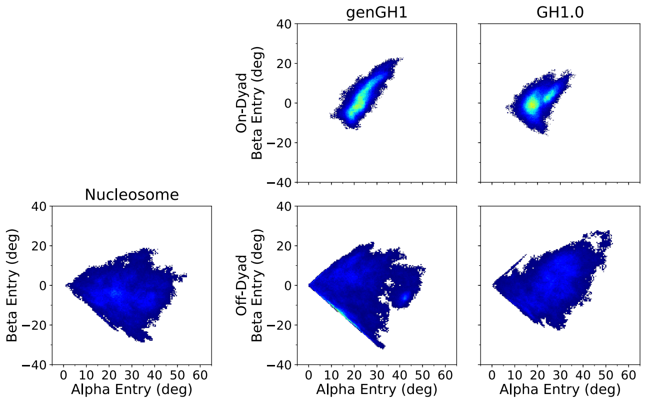


Figure S16: Density plots of entry linker DNA angles. Each plot shows a 2D histogram of the alpha versus the beta entry linker DNA angles. Density ranges from dark blue (lowest) to red (highest). Each plot contains the linker DNA from the nucleosome core particle along with genGH1 and GH1.0 in the on- and off-dyad binding modes.

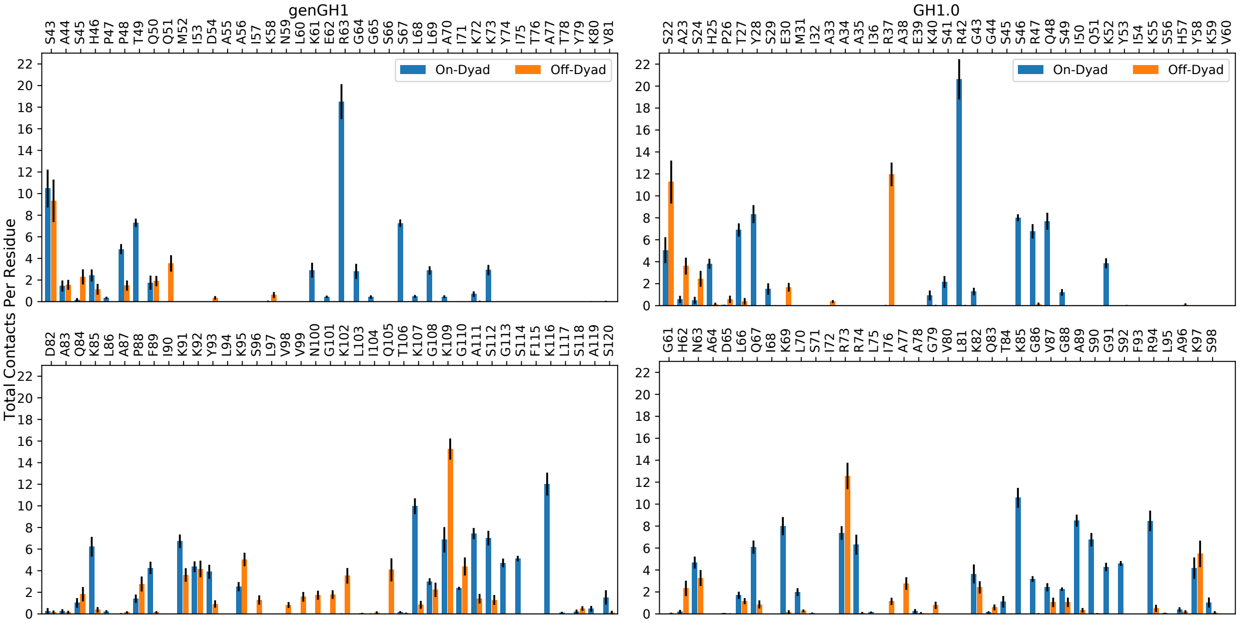


Figure S17: Contacts of genGH1 and GH1.0 residues with DNA in both the on- (blue) and off-dyad (orange) binding modes.

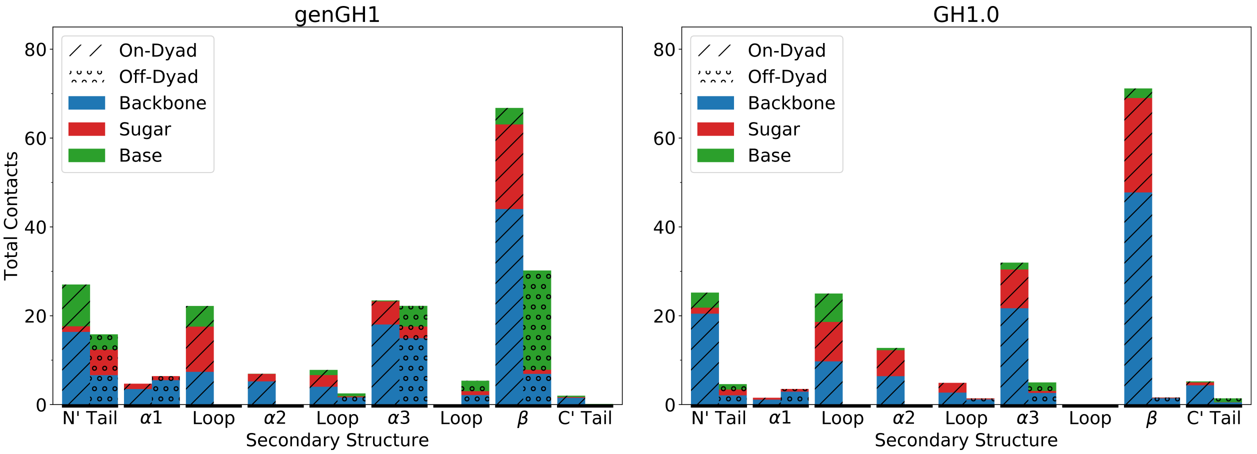


Figure S18: Total contacts of genGH1 and GH1.0 secondary structures with DNA in both the on- and off-dyad binding modes. Each bar is shown as a percentage of contacts with the backbone (blue), sugar (red), and base (green) of the DNA.

### S5 GH1.0 Pentamutant

#### S5.1 System Construction

The GH1.0 pentamutant structure was constructed using the previously described GH1.0 wild type structures as a template (see Methods in the main text). The following mutations were introduced: R47L, R74S, V80K, V87K, and K97A. These mutations were applied to the structure via the *tleap* module of Amber18.

#### S5.2 Molecular Dynamics Simulations and Analysis Methods

With the exception of the system construction described above, simulations and analyses of the GH1.0 pentamutant were all conducted in the same manner as detailed in the Methods section of the main text.

#### S5.3 Tables

Table S4: Energy differences (kcal/mol) between on and off-dyad binding states as estimated by MM/GBSA analysis. A negative value indicates more favorable binding in the on-dyad state.

| Linker Histone | Binding Mode | $\Delta E_{total}$ | $\Delta E_{internal}$ | $\Delta E_{elec}$ | $\Delta E_{VdW}$ | $\Delta\Delta E_{total}$ | $\Delta\Delta E_{internal}$ | $\Delta\Delta E_{elec}$ | $\Delta\Delta E_{VdW}$ |
|---|---|---|---|---|---|---|---|---|---|
| GH1.0-pMut | On-Dyad | -98.2 ± 40.1 | 5.4 ± 32.8 | 62.3 ± 27.3 | -165.8 ± 22.2 | -39.0 ± 39.3 | -4.2 ± 33.3 | 35.6 ± 25.2 | -70.3 ± 22.8 |
| | Off-Dyad | -59.2 ± 37.0 | 9.6 ± 32.8 | 26.7 ± 23.9 | -95.5 ± 22.3 | | | | |

#### S5.4 Figures

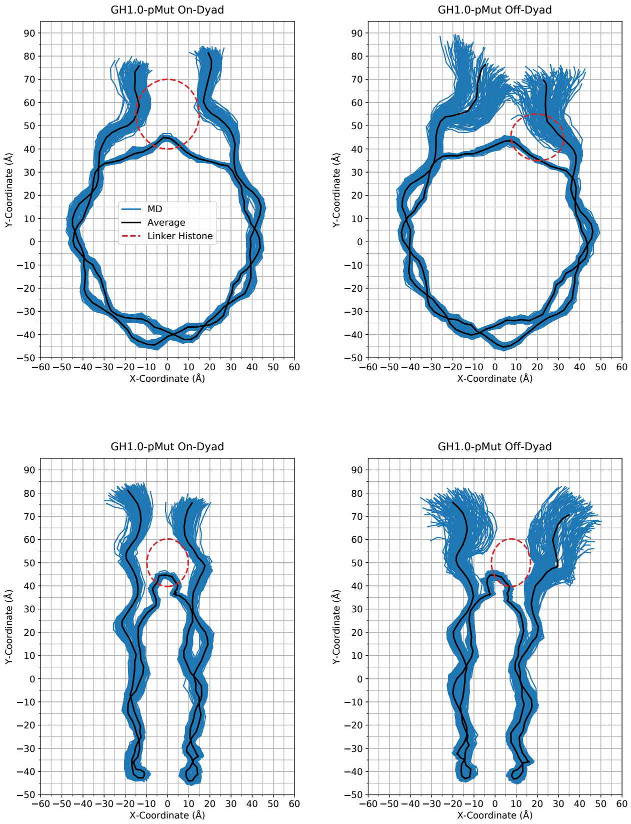


Figure S19: In-plane (top) and out-of-plane (bottom) DNA motions sampled by the GH1.0 pentamutant in the on- and off-dyad binding modes. Shown in blue are configurations sampled throughout the MD simulation (150 representative frames) while the average configuration is shown in black. For reference, the approximate position of the linker histone is shown as a dashed-line red ellipse. Figures inspired by work from Shaytan *et al.*[93]

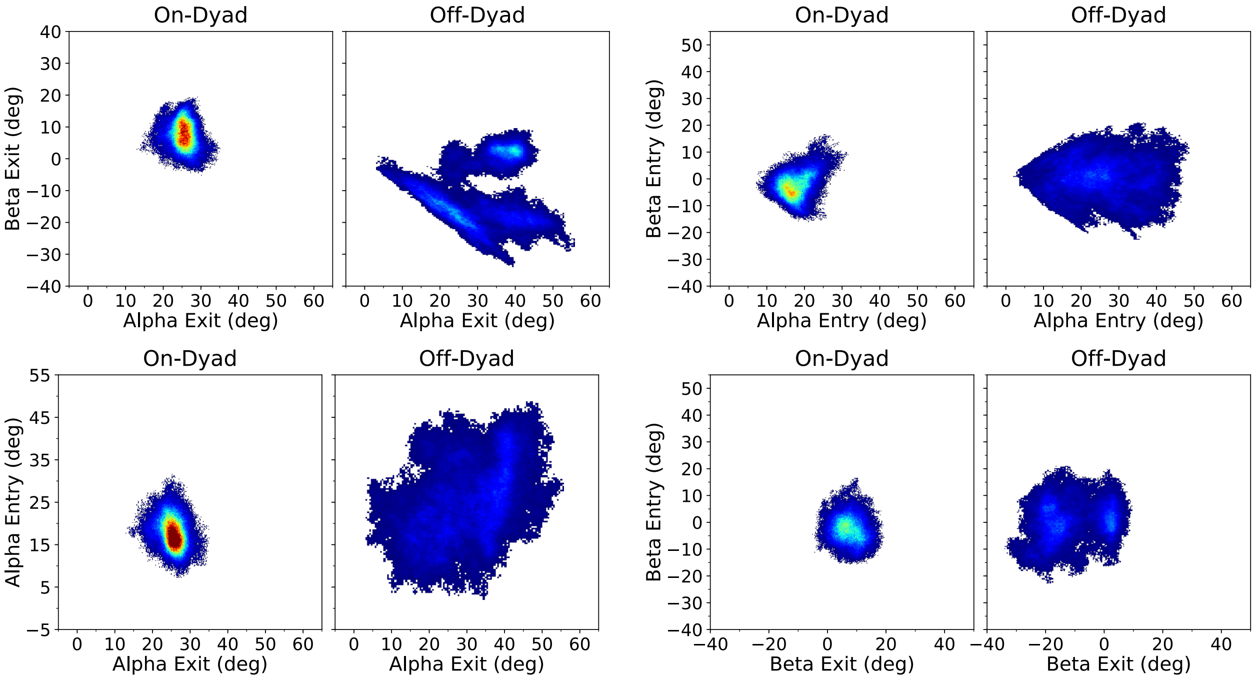


Figure S20: Density plots of linker DNA α and β angles for exit and entry DNA from the on- and off-dyad GH1.0 pentamutant systems. Densities range from dark blue (lowest) to red (highest).

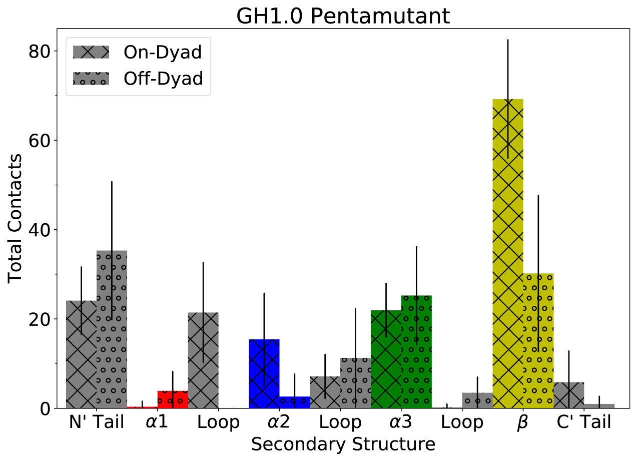


Figure S21: Total contacts of the GH1.0 pentamutant secondary structures with DNA in both the on- and off-dyad binding modes. The secondary structure is broken down into the following: alpha helix α1 (red), alpha helix α2 (blue), alpha helix α3 (green), β-sheet (yellow), N-terminal tail (grey), C-terminal tail (grey), and three loops (grey).

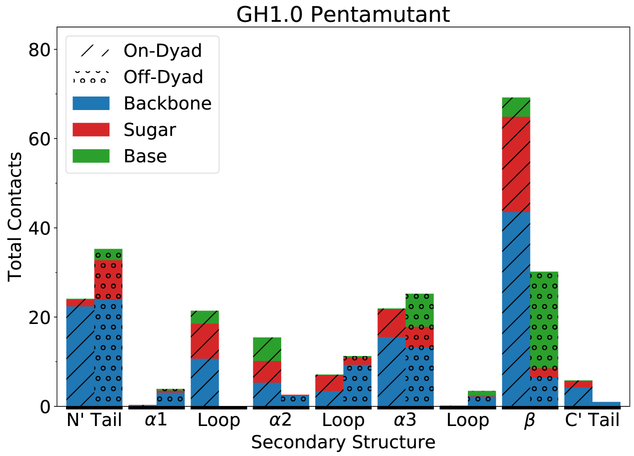


Figure S22: Total contacts of the GH1.0 pentamutant secondary structures with DNA in both the on- and off-dyad binding modes. Each bar is shown as a percentage of contacts with the backbone (blue), sugar (red), and base (green) of the DNA.

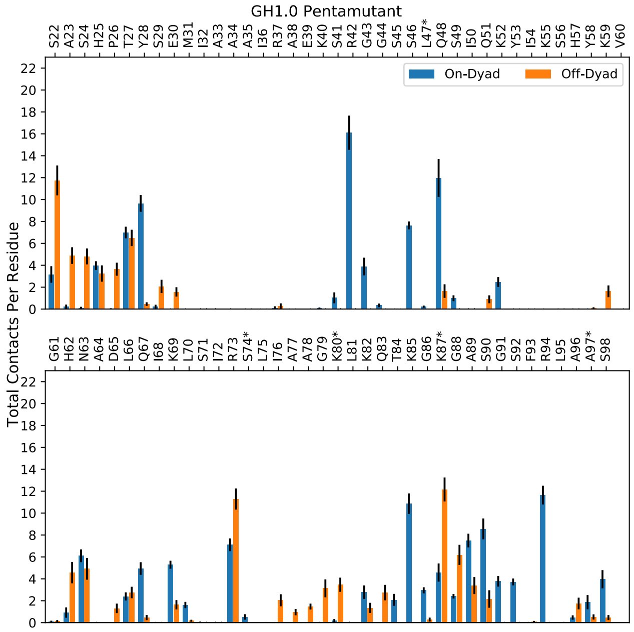


Figure S23: Contacts of the GH1.0 pentamutant residues with DNA in both the on- (blue) and off-dyad (orange) binding modes. Residues marked with a '*' were mutated from the wild type GH1.0 to the analogous genGH1 residues (see System Construction above in Section S5.1).
